# Genetically Labeled Premyelinating Oligodendrocytes: Bridging Oligodendrogenesis and Neuronal Activity

**DOI:** 10.1101/2024.12.27.630559

**Authors:** Aksheev Bhambri, Phu Thai, Songtao Wei, Han-Gyu Bae, Daniela Barbosa, Tripti Sharma, Ze Yu, Chao Xing, Jun Hee Kim, Guoqiang Yu, Lu O. Sun

## Abstract

To myelinate axons, oligodendrocyte precursor cells (OPCs) must stop dividing and differentiate into premyelinating oligodendrocytes (preOLs). PreOLs are thought to survey and begin ensheathing nearby axons, and their maturation is often stalled at human demyelinating lesions. Lack of genetic tools to visualize and manipulate preOLs has left this critical differentiation stage woefully understudied. Here, we generated a knock-in mouse line that specifically labels preOLs across the central nervous system. Genetically labeled preOLs exhibit distinct morphology, unique transcriptomic and electrophysiological features, and do not overlap with OPCs. PreOL lineage tracing revealed that subsets of them undergo prolonged maturation and that different brain regions initiate oligodendrogenesis with the spatiotemporal specificity. Lastly, by fate mapping preOLs under sensory deprivation, we find that neuronal activity functions within a narrow time window of preOL maturation to promote their survival and successful integration. Our work provides a new tool to probe this critical cell stage during axon ensheathment, allowing for fine dissection of axon-oligodendrocyte interactions.

## INTRODUCTION

In the central nervous system (CNS), myelin is produced by oligodendrocytes to provide support and insulation for axons.^1^ To become myelinating oligodendrocytes, oligodendrocyte progenitor cells (OPCs) must exit cell cycle and differentiate to premyelinating oligodendrocyte (preOLs).^2^ PreOLs are long thought to be a transient cell stage, existing in the adult mouse brain for approximately 1.8 days.^3^ Depending on the brain regions, a proportion or all of them (20% - 100%) undergo programmed cell death prior to myelination during development and into adulthood.^3–7^ Despite being transient and vulnerable, preOLs are at a critical stage of myelination, known as axon ensheathment. At this stage, they extend several processes to survey and begin ensheathing nearby axons.^8,9^ Due to their critical roles in oligodendrogenesis, dysregulation of preOL maturation is observed in demyelinating diseases such as multiple sclerosis and perinatal white matter injury (pWMI).^10,11^

Despite their importance in health and disease, preOLs still remain poorly characterized. This is partly due to the dearth of methods to label and distinguish them from other oligodendrocyte linage cells, as well as the lack of tools to lineage trace and manipulate them. For instance, preOLs can be visualized *in vivo* by antibodies raised against BCAS1 and DM20.^4,12^ However, both antibodies show extended expression in mature oligodendrocytes. *Enpp6* and *LncOL1* have been recently identified as two preOL markers.^13–15^ Since both are RNA markers, their usages are largely constrained to instantaneous visualization of preOL cell bodies, making it impossible to delineate the entire morphology of preOLs and to fate map them during myelination. Several genetic tools, such as *Cnp-Cre* and *Plp-CreER* mouse lines, have been utilized to label and manipulate genes starting from the preOL stage.^16,17^ Nevertheless, they also exhibit extended expression in mature oligodendrocytes and can leak into other cell types, making it challenging to specifically probe preOLs.^18–20^ Lastly, multi-photon live imaging studies have provided key information during oligodendrogenesis, including the preOL stage.^3,21,22^ However, due to the lack of a direct reporter, preOLs are visualized indirectly through the disappearance of a fluorescent marker that labels OPC,^3^ which may still be present in the preOLs despite their degradation.

In humans, oligodendrogenesis and neuronal activity are coupled together through reciprocal interactions during development and under diseased conditions.^23^ Recent studies in animal models, using complex motor learning,^13,24^ fear conditioning,^25^ pharmacogenetic manipulation,^26^ and optogenetic stimulation^27^, have shown that neuronal activity can significantly promote oligodendrogenesis and myelination. This process, known as activity-dependent myelination, has been extensively studied in the context of OPC proliferation and subsequent differentiation.^3,13,24–30^ Meanwhile, it has been long hypothesized that neuronal activity can directly act on existing preOLs to modulate myelination.^31^ A study using a whisker trimming paradigm showed that decreased sensory input reduces mature oligodendrocyte number during development, possibly through reduced immature oligodendrocyte survival.^32^ On the other hand, a recent live-animal imaging study revealed that sensory enrichment increases the integration of newly formed oligodendrocytes into the neuronal circuits.^3^ Due to lack of specific genetic reagents, direct evidence that neuronal activity regulates preOL survival and integration, the bottleneck during axon ensheathment, is still lacking.

To fulfil the need for a specific *in vivo* tool, we developed a preOL-specific driver mouse line, the *Enpp6-IRES-CreER^T2^*, allowing visualization and lineage tracing of preOLs throughout the CNS. Further characterization shows that genetically labeled preOLs are postmitotic and do not overlap with OPCs. Through genetic labeling, we demonstrate the unique histological, transcriptomic, and electrophysiological features of preOLs, as well as their spatiotemporal specificity in integrating into diverse brain regions. The transcriptomic dataset for genetically labeled preOLs has been made available on a user-friendly website (utswbioinformatics.shinyapps.io/scrna_preol/). Finally, using this preOL driver line, we show that neuronal activity acts directly on existing preOLs within a critical time window to promote their survival and integration. Together, our work provides a new tool to genetically probe this critical rate-limiting stage during oligodendrogenesis and myelination, allowing refined dissection of oligodendrocyte-axon interactions in health and disease.

## RESULTS

### Generation of a knock-in mouse line to genetically probe premyelinating oligodendrocytes (preOLs) across the central nervous system

To genetically label premyelinating oligodendrocytes (preOLs), we took advantage of *Enpp6*, a well-known marker for preOLs (Figure 1A),^13^ to create a *Enpp6-IRES-CreER^T2^* driver mouse line (hereafter referred to as *Enpp6-CreER^T2^*; Table S1). To prevent disruption of *Enpp6* gene expression, we utilized the *Easi*-CRISPR method to insert *IRES-CreER^T2^* in the 3’untranslated region (UTR) of the gene, a few bases downstream of the stop codon in the *Enpp6* coding sequence (Figure 1B). To visualize *Enpp6*^+^ cells, the *Enpp6-CreER^T2^* animals were crossed with the Ai9 reporter mice, and the *Enpp6-CreER^T2^; Ai9/+* pups were administered with 4-hydroxytamoxifen (4HT) at postnatal day 8 (P8). At 1 day post injection (1 dpi), *Enpp6-CreER^T2^; Ai9/+* sagittal brain sections readily showed numerous DsRed^+^ cells present throughout the brain (Figure 1C). At higher magnification, the DsRed^+^ cells exhibited extensive arborization and a ramified morphology, which is typically observed in preOLs (Figure 1D).^4^ Importantly, DsRed^+^ cells were not found in *Enpp6-CreER^T2^; Ai9/+* mice after oil injection (Figures S1A and S1A’), indicating that there is no 4HT-independent recombination with this driver line.

**Figure 1.**
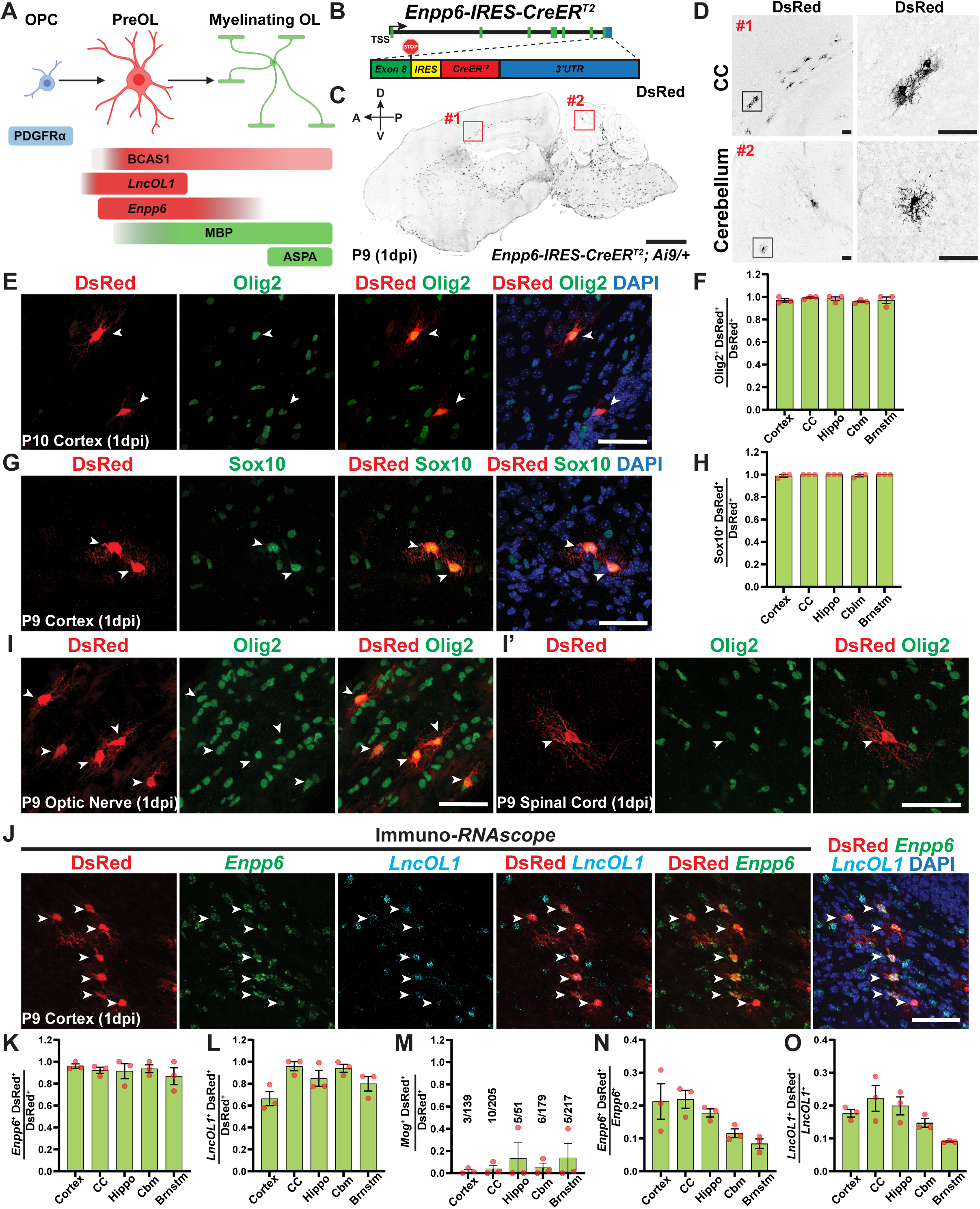
Design and generation of a knock-in mouse line to genetically label premyelinating oligodendrocytes (preOLs) throughout the central nervous system. **(A)** Schematic showing markers used to label and visualize oligodendrocyte progenitor cells (OPCs), premyelinating oligodendrocytes (preOLs), and myelinating oligodendrocytes (myelinating OLs). **(B)** Schematic showing the design of *Enpp6-IRES-CreER^T2^* driver line. The *IRES-CreER^T2^* cassette was inserted into the 3’UTR of *Enpp6*, immediately after the stop codon in exon 8. **(C)** Sagittal brain section of postnatal day 9 (P9) *Enpp6-CreER^T2^; Ai9/+* mouse 1 day post injection (dpi) with 150 mg/kg 4-hydroxytamoxifen (4HT). The DsRed^+^ cells were visualized by immunohistochemistry using an anti-DsRed antibody. The red insets mark the regions selected for high-magnification images at corpus callosum (#1) and cerebellum (#2). **(D)** High magnification images of #1 and #2 insets in (C) showing the ramified, “spider-like” morphology distinctly observed in preOLs. **(E)** Representative images of P9 *Enpp6-CreER^T2^; Ai9/+* cortex administered with 150 mg/kg 4HT at 1 dpi using antibodies raised against DsRed (red) and Olig2 (green). **(F)** Quantifications of the proportion of DsRed^+^ Olig2^+^ cells over the total number of DsRed^+^ cells in P9 *Enpp6-CreER^T2^; Ai9/+* cortices. **(G and H)** Immunohistochemistry (G) and quantification (H) of P9 *Enpp6-CreER^T2^; Ai9/+* brain sections stained with DsRed (red) and Sox10 (green) antibodies. **(I and I’)** Representative confocal images of P9 *Enpp6-CreER^T2^; Ai9/+* optic nerve (I) and spinal cord (I’) stained with DsRed (red) and Olig2 (green) antibodies. **(J)** Immunohistochemistry combined with RNAScope using DsRed antibody (red), *Enpp6* probe (green), and *LncOL1* probe (cyan) in P9 *Enpp6-CreER^T2^; Ai9/+* cortex. **(K and L)** Quantifications of the proportion of *Enpp6*^+^ DsRed^+^ cells (K) and *LncOL1*^+^ DsRed^+^ cells (L) among the total DsRed^+^ cells. **(M)** Quantification of *Mog*^+^ DsRed^+^ cell proportion among the total DsRed^+^ cells. **(N and O)** Quantifications of the proportions of *Enpp6*^+^ DsRed^+^ and *LncOL1*^+^ DsRed^+^ cells among the total *Enpp6*^+^ cells (N) and *LncOL1*^+^ cells (O), respectively. Error bars represent SEM. Scale bars: 1 mm in (C); 50 µm in (D), (E), (G), (I), (I’), and (J). Close circles in (F), (H), and (K)-(O) represent individual animals. n=3 animals per category. CC: Corpus Callosum; Hippo: Hippocampus; Cblm: Cerebellum; Brnstm: Brainstem.

To characterize *Enpp6-CreER^T2^*-labeled cells, we co-stained the DsRed^+^ cells with Olig2 and Sox10, two pan-oligo lineage markers, and quantified the proportion of co-labeled cells among the total DsRed^+^ cells in the brain (Figures 1E and 1G). Nearly all DsRed^+^ cells were Olig2^+^ (97.48% ± 1.6%; Figure 1F) and So×10^+^ (99.58% ± 0.36%; Figure 1H). Moreover, *Enpp6-CreER^T2^*exhibited no or minimal labeling in astrocytes, microglia, and neurons (Figures S1C-E’). To determine if the *Enpp6-CreER^T2^* could also label oligodendrocyte lineage cells in other central nervous system (CNS) tissues, we analyzed optic nerves and spinal cords from P9 *Enpp6-CreER^T2^; Ai9/+* pups at 1 dpi. Both the optic nerve and spinal cord showed ramified DsRed^+^ cells that were also Olig2^+^ (Figure 1I, I’). By contrast, sciatic nerves from the peripheral nervous system (PNS) exhibited no DsRed^+^ cell labeling (Figure S1B).

To test if *Enpp6-CreER^T2^* selectively labels preOLs, we performed immunohistochemistry combined with *in situ* hybridization. We used a DsRed antibody to visualize genetically labeled cells and *in situ* probes, such as *Enpp6* and *LncOL1,* for preOLs (Figure 1J). We observed that nearly all DsRed^+^ cells were *Enpp6^+^* (Figure 1K), while ∼84.76% of DsRed^+^ cells were *LncOL1^+^* (Figure 1L). Moreover, only a small portion of DsRed^+^ cells are *Mog^+^*(Figure 1M), a marker for mature oligodendrocytes, suggesting that the majority of DsRed^+^ cells are still at the immature, premyelinating stage (see also Figures 2 and 3). We calculated the labeling efficiency of the *Enpp6-CreER^T2^* line and found that approximately 16.52% of total *Enpp6^+^* cells and approximately 16.57% of total *LncOL1^+^* cells were DsRed^+^ on average among diverse brain regions at P9 (Figures 1N and 1O). Thus, the *Enpp6-CreER^T2^* knock-in mouse line demarcates subsets of preOLs across the CNS.

**Figure 2.**
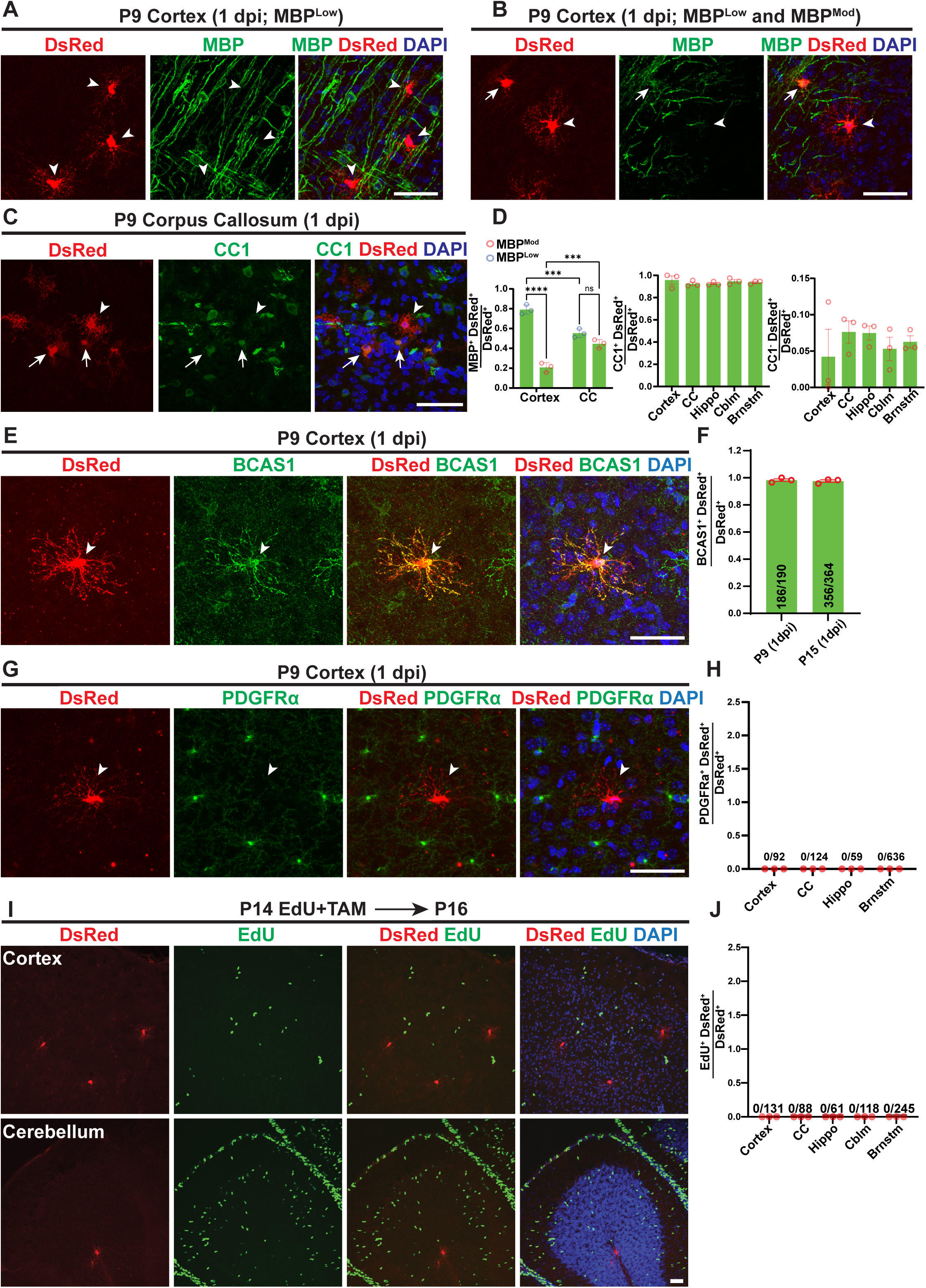
Genetically labeled preOLs are postmitotic and uncover a hidden stage after OPCs initiate differentiation. **(A and B)** Representative images of P9 *Enpp6-CreER^T2^; Ai9/+* sagittal brain sections at 1 day post injection (dpi) stained by DsRed (red) and myelin basic protein (MBP; green) antibodies. Arrowheads indicate DsRed^+^ MBP^Low^ cells in (A). The arrowhead indicates a DsRed^+^ MBP^Low^ cell while the arrow shows a DsRed^+^ MBP^Moderate^ cell in (B). **(C)** Representative images of P9 *Enpp6-CreER^T2^; Ai9/+* sagittal brain sections at 1 dpi stained by DsRed (red) and CC1 (green) antibodies. The arrowhead indicates a DsRed^+^ CC1^-^ cell while arrows show DsRed^+^ CC1^+^ cells. Quantification of MBP^Low^ DsRed^+^ and MBP^Mod^ DsRed^+^ cells among the total DsRed^+^ cells (left column) and quantification of CC1^+^ DsRed^+^ cells and CC1^-^ DsRed^+^ cells among the total DsRed^+^ cells (right two columns). **(E)** Representative images of P9 *Enpp6-CreER^T2^; Ai9/+* sagittal brain sections immunostained by DsRed (red) and BCAS1 (green) antibodies at 1 dpi. **(F)** Quantifications of BCAS1^+^ DsRed^+^ cell proportion among the total DsRed^+^ cells at P9 and P15. **(G and H)** Representative images (G) and quantification (H) of PDGFRα^+^ DsRed^+^ cell proportion among the total DsRed^+^ in P9 *Enpp6-CreER^T2^; Ai9/+* sagittal brain sections at 1 dpi immunostained by DsRed (red) and PDGFRα (green) antibodies. **(I and J)** Representative images (I) and quantification (J) of EdU^+^ (green) DsRed^+^ cell proportion among the total DsRed^+^ cells (red) in P16 *Enpp6-CreER^T2^; Ai9/+* brains at 2 dpi of tamoxifen. Error bars represent SEM. Scale bars: 50 µm. Circles in (D), (F), (H), and (J) represent individual animals. n=3 animals per category. The numbers at each bar for (F), (H), and (J) indicate the number of subsets of DsRed^+^ cells relative to the total number of DsRed^+^ cells quantified. CC: Corpus Callosum; Hippo: Hippocampus; Cblm: Cerebellum; Brnstm: Brainstem.

**Figure 3.**
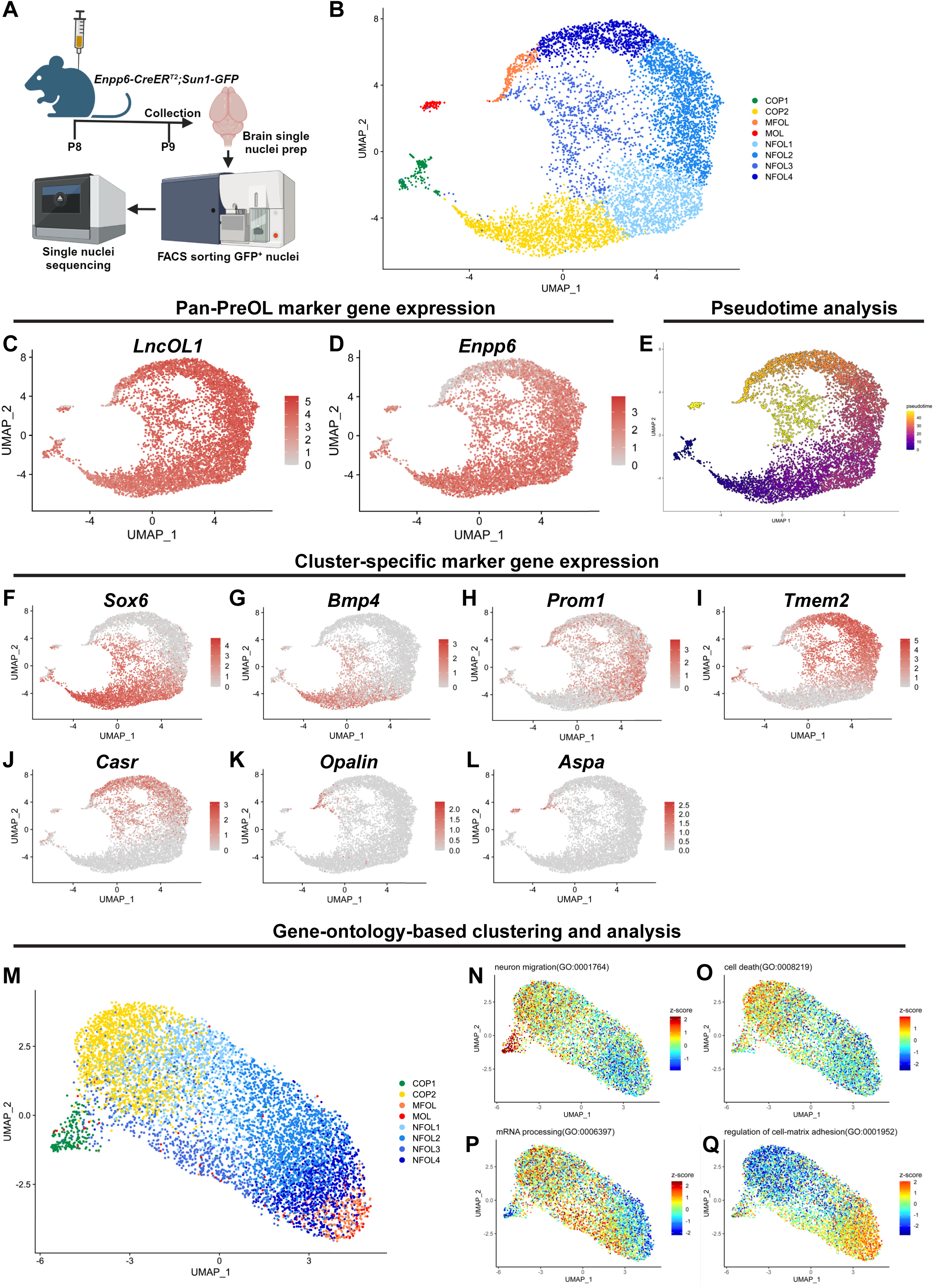
Single nuclei RNA sequencing analysis of genetically labeled preOLs reveals distinct marker gene expression and differentiation-stage-dependent biological processes related to preOL survival and axon ensheathment. **(A)** Schematic showing the single nuclei RNA sequencing (snRNAseq) of genetically labeled preOLs (*Enpp6-CreER^T2^; Sun1-GFP/+*). **(B)** Uniform Manifold Approximation and Projection (UMAP) plots showing the preOL lineage composition using Seurat-based clustering. **(C and D)** UMAP showing the expression of pan-preOL markers, *Enpp6* (C) and *LncOL1* (D), in genetically labeled preOLs. **(E)** Pseudo-time analysis using Seurat package shown as UMAP indicates the developmental trajectory of genetically labeled preOLs. More differentiated cells are shown in yellow while younger or immature preOLs are shown in blue. **(F-L)** UMAPs showing the expression of cluster-enriched markers including *Sox6*, *Bmp4*, *Prom1*, *Tmem2*, *Casr*, *Opalin*, and *Aspa* across preOL lineage. **(M)** UMAP plots showing gene-ontology (GO)-based clustering of genetically labeled preOLs. **(N-Q)** UMAPs showing the z-scores for GO-terms: “neuron migration” (N), “cell death” (O), “mRNA processing” (P), and “cell matrix regulation” (Q).

### Genetically labeled preOLs are post-mitotic and reveal a hidden stage after OPCs initiate differentiation

PreOLs are thought to be at a transient differentiation stage when OPCs cease proliferation and begin expressing myelin proteins.^3^ We next characterized the genetically labeled preOLs (P9 *Enpp6-CreER^T2^; Ai9/+,* 1 dpi) with known markers for postmitotic oligodendrocytes and OPCs. We found that the majority of DsRed^+^ cells in the cortex exhibited a ramified morphology and showed either no or very low MBP expression (arrowheads in Figure 2A; quantified in Figure 2D, 93/118 cells from 3 animals), whereas a small fraction of them began expressing MBP and elaborating longitudinally formed processes (arrows in Figure 2B; quantified in Figure 2D, 25/118 cells from 3 animals). By contrast, ∼40% of DsRed^+^ cells in the corpus callosum exhibit moderate MBP expression in their cell bodies (quantified in Figure D). Intriguingly, while majority of the DsRed^+^ cells were CC1^+^, a marker for postmitotic oligodendrocytes,^33,34^ approximately 8% of the DsRed^+^ cells showed no CC1 expression across diverse brains regions (Figures 2C and 2D).

We suspect *Enpp6-CreER^T2^* expression begins in an as-yet-unstudied early phase of preOL maturation, during which MBP and CC1 remain unexpressed. To test it, we stained the genetically labeled preOLs with BCAS1, a preOL marker.^12^ The majority of the DsRed^+^ cells showed BCAS1 positivity in P9 and P15 *Enpp6-CreER^T2^; Ai9/+* mouse brain sections at 1 dpi (Figures 2E and 2F), confirming that the genetically labeled cells are indeed preOLs. Furthermore, the DsRed^+^ cells are not at OPC stage as none of them expressed PDGFRα, an OPC marker (Figures 2G and 2H).^35^ To rule out the possibility that *Enpp6-CreER^T2^*could label a small population of proliferating OPCs that readily downregulate PDGFRα expression, we performed 5-ethynyl-2’-deoxyuridine (EdU) injection in P14 *Enpp6-CreER^T2^; Ai9/+* animals that were simultaneously administered with tamoxifen. We did not observe any DsRed^+^ EdU^+^ cells at 2 dpi across diverse brain regions (Figures 2I and 2J). Therefore, *Enpp6-CreER^T2^* does not label OPCs but delineates an early stage in preOL maturation.

### Genetically labeled preOLs express distinct marker genes and exhibit differentiation-stage-dependent biological processes

Our characterization of *Enpp6-CreER^T2^* suggests that it labels the earliest stage after OPCs exit cell cycle, known as committed oligodendrocyte precursor (COP), that remains poorly understood.^36,37^ To characterize it and further determine transcriptomic heterogeneity among *Enpp6-CreER^T2^*-labeled cells, we crossed the *Enpp6-CreER^T2^* animals with the nucleus reporter line, *Sun1-GFP,*^38^ and purified GFP^+^ nuclei 1 dpi of 4HT for single nuclei RNA sequencing (snRNA-seq) (Figure 3A). After quality control assessment and filtering, we obtained 8,493 nuclei from two mouse brains. These nuclei were analyzed using Seurat package, and gene expression matrix was used for unsupervised clustering (Figure S2A). After excluding contaminating clusters enriched with choroid plexus markers such as *Ttr* (Figure S2F),^39^ and those enriched with blood cell markers (Figure S2G), we obtained 8 major clusters. These clusters were annotated into COP (COP1-2), newly formed oligodendrocyte (NFOL1-4), myelin forming oligodendrocyte (MFOL), and mature oligodendrocyte (MOL; Figure 3B). These clusters exhibited high similarity scores to previously reported COP, NFOL, MOL, and MFOL clusters, respectively (Figures S2B-E; Table S2).^36^

As expected, nuclei in all the clusters showed high expression levels of pan-preOL genes, such as *Enpp6* and *LncOL1* (Figures 3C and 3D). Moreover, the known preOL-enriched genes exhibited moderate-to-high expression throughout the genetically labeled preOL lineage (Figures S2H-P), further confirming that our dataset primarily consists of preOLs. Our analysis unveiled cluster-enriched expression of preOL markers, such as *Sox6* (COP1-2, NFOL1-2; Figure 3F), *Bmp4* (COP2 and NFOL1; Figure 3G), *Prom1* (NFOL1-4; Figure 3H), and *Tmem2* and *Casr* (NFOL2-4; Figures 3I and 3J). Intriguingly, we identified an early stage of COP, termed COP1, which likely represents cells just exiting the OPC cell cycle. COP1 expressed a set of marker genes distinct from COP2 (Figure S3; see also Table S3) and was identified as the earliest preOL stage by pseudotime analysis (Figure 3E). We have made this dataset easily accessible as a user-friendly website (utswbioinformatics.shinyapps.io/scrna_preol/).

To begin understanding the stage-dependent function through preOL maturation, we performed gene ontology (GO)-based clustering and analysis (Figure 3M). GO-based clustering showed limited heterogeneity among the nuclei, indicating a remarkable similarity between individual nuclei. Despite limited diversity, we identified several cluster-enriched biological processes (Figures 3N-Q). For instance, the GO term “neuron migration” was highly enriched in COP1 (Figure 3N), while the GO term “cell death” was enriched in COP2 (Figure 3O), likely representing two key chronological steps that occur during early preOL maturation.^3,22^ We also found that NFOLs exhibit enriched GO terms such as “mRNA processing” (Figure 3P), steroid biosynthesis (Figure S2Q), lipid transport (Figure S2R), and cholesterol metabolism (Figure S2S). Lastly, the GO term “cell adhesion” was enriched in MFOL and MOLs (Figure 3Q), together suggesting the sequential molecular and cellular events required for myelin protein synthesis and axon ensheathment during preOL lineage progression.

### Genetically labeled preOLs emerge and integrate into diverse brain regions with spatiotemporal specificity

Oligodendrogenesis and myelination peak early postnatally and continue throughout animals’ lifespan.^21^ To determine the spatiotemporal pattern of oligodendrogenesis in the developing murine brain, we characterized the genetically labeled preOLs across diverse brain regions at multiple time points. We administered 4HT to *Enpp6-CreER^T2^; Ai9/+* mice at P8, P14, and P21, and analyzed the brains at 1 dpi. At P9, we observed that the white matter regions of cerebellum and brainstem had significantly higher DsRed^+^ preOL densities compared to grey matter regions such as hippocampus and cortex (Figure 4A; quantified in Figure 4B). Moreover, the corpus collosum continued to exhibit the highest DsRed^+^ preOL density at P15 and P22 (Figures 4A’-A”; quantified in Figures 4C and 4D), whereas hippocampus was among the latest brain regions to harbor preOLs (Figure 4H). These results were further confirmed by analyzing the total number of *LncOL1^+^* preOLs and their spatiotemporal patterns (Figures 4E-G). We noticed that the labeling efficiency of *Enpp6-CreER^T2^* gradually decreased as the brain developed (Figure S4K-O). Despite reduced efficiency, *Enpp6-CreER^T2^* did not exhibit preferential preOL labeling across brain areas at any of the given time points (Figures S4A-J).

**Figure 4.**
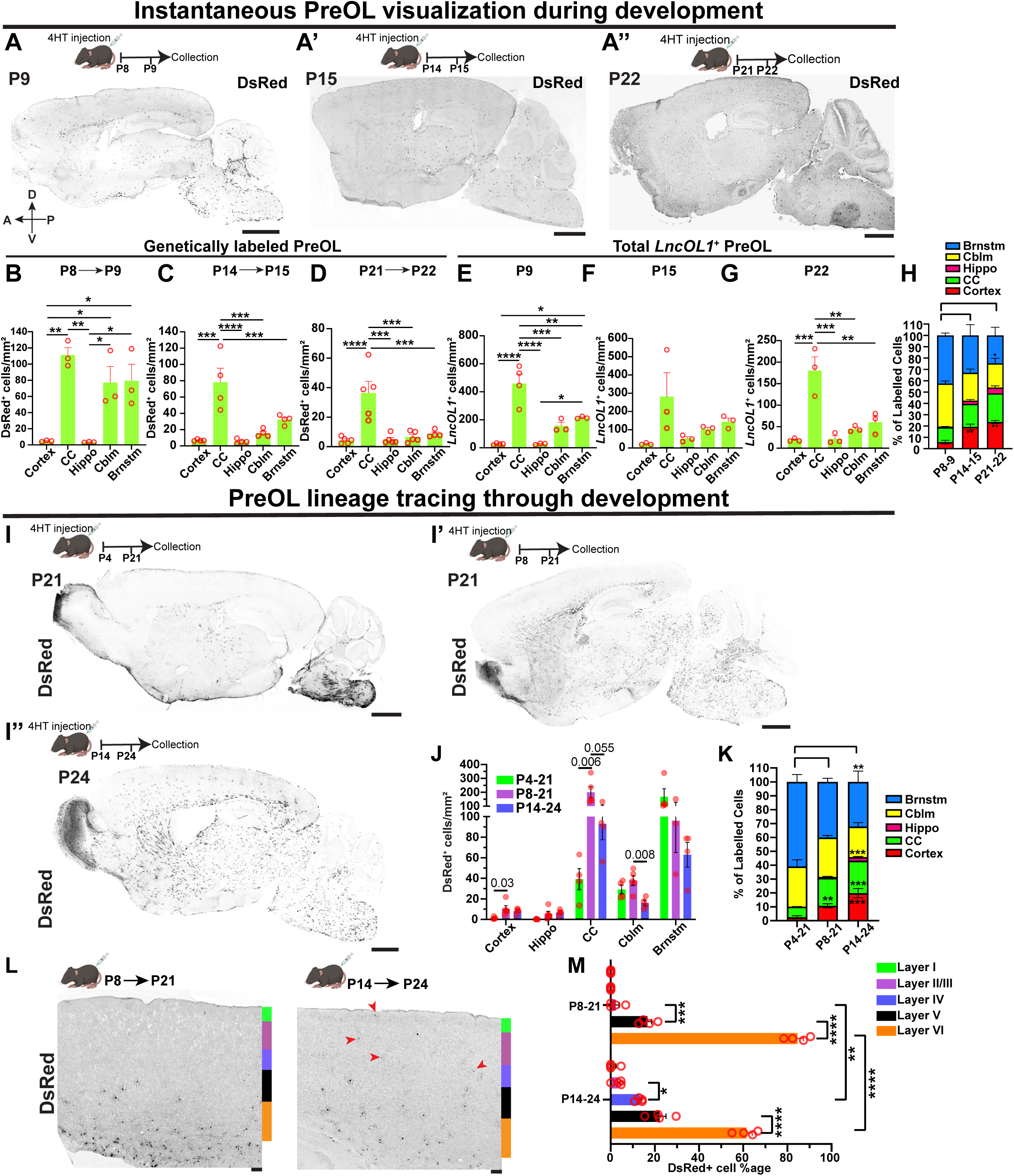
Genetically labeled preOLs emerge and integrate into diverse brain regions spatiotemporally. **(A-A”)** Representative images of *Enpp6-CreER^T2^; Ai9/+* sagittal brain sections at P9 (A), P15 (A’) and P21 (A”), 1 day post injection (dpi) of 150 mg/kg 4-hydroxytamoxifen (4HT). DsRed^+^ cells were visualized using an antibody raised against DsRed. **(B-D)** Quantification of DsRed^+^ cell density 1 dpi on P9 (B, n=3), P15 (C, n=4), and P22 (D, n=5) sagittal brain sections. **(E-G)** Quantification of *LncOL1*^+^ cell density on P9 (E), P15 (F), and P22 (G) sagittal brain sections. **(H)** Percentage of DsRed^+^ cells plotted across brain regions on P9, P15, and P22 *Enpp6-CreER^T2^; Ai9/+* sagittal brain sections. **(I-I”)** Representative images of P21 or P24 *Enpp6-CreER^T2^; Ai9/+* sagittal brain sections administered with 100 mg/kg 4HT at P4 (I), P8 (I’), and P14 (I”). DsRed^+^ cells were visualized by an antibody raised against DsRed. **(J)** Quantification of DsRed^+^ cell density across brain regions where preOLs were lineage-traced at P4, P8, and P14. n=4. **(K)** Percentage of DsRed^+^ cells plotted across *Enpp6-CreER^T2^; Ai9/+* brain regions where preOLs were lineage-traced at P4, P8 and P14. **(L)** Representative images showing DsRed^+^ cell in the somatosensory cortex of P21 (left) and P24 (right) *Enpp6-CreER^T2^; Ai9/+ sagittal* sections that were lineage-traced from P8 and P14, respectively. Red arrowheads mark DsRed^+^ cells observed in the cortical layers I-IV. **(M)** Quantification of DsRed^+^ cell percentage across cortical layers where preOLs were lineage-traced from P8 to P21 and from P14 to P24. Error bars represent SEM. Circles in (B)-(G), (J), and (M) represent individual animals. One-way ANOVA followed by Tukey’s multiple comparisons test for (B)-(H), (J), (K), and (M). *p<0.05, **p<0.01, ***p<0.001, and ****p<0.0001. Scale bars: 1 mm in (A)-(A”); 1 mm in (I)-(I”); and 100 µm in (L). CC: Corpus Callosum; Hippo: Hippocampus; Cblm: Cerebellum and Brnstm: Brainstem.

To understand how preOLs mature and incorporate into neuronal circuits over time, we performed lineage tracing experiments at three developmental stages in *Enpp6-CreER^T2^; Ai9/+* animals. The mice were injected at P4, P8, or P14 and analyzed around the weaning age (P21 or P24). Oligodendrocytes originating from preOLs formed at P4 were predominantly located in the brainstem and cerebellum (Figure 4I). In contrast, preOLs formed at P8 and lineage-traced occupied many other brain regions, such as the corpus collosum, striatum, midbrain, and deep cortical layers (Figures 4I’ and 4L; quantified in Figures 4J, 4K, and 4M). Finally, oligodendrocytes lineage-traced from P14 preOLs began to appear in the hippocampus and upper cortical layers (Figures 4I” and 4L; quantified in Figures 4J, 4K, and 4M), representing two brain areas with a late onset of myelination. Therefore, preOLs emerge and integrate into diverse brain regions with spatiotemporal specificity during development.

### Subsets of genetically labeled preOLs undergo prolonged maturation during early brain development

PreOLs are considered to be at a transition stage during oligodendrogenesis.^3^ To determine how transient they are, we performed lineage tracing on P8 *Enpp6-CreER^T2^; Ai9/+* animals and analyzed genetically labeled preOLs throughout their maturation at 2, 4, 6, 8, and 13 dpi (Figure 5A). We examined the DsRed^+^ cells in the cortex and categorized them as “ramified” or “sheath-forming” cells based on their morphology (Figure 5B). The majority of DsRed^+^ cells exhibited a “sheath-forming” morphology by 6 dpi. Surprisingly, approximately 3% of total DsRed^+^ cells continued to exhibit ramified morphology up to 8 dpi in the cortex (Figures 5C and Figures S5; quantified in Figure 5D). These ramified cells were randomly distributed in both upper and deep cortical layers (Figures S5A and S5C). They were indeed preOLs, as evidenced by their low MBP protein expression and BCAS1^+^ status (Figures 5C and S5). By contrast, although we observed abundant ramified DsRed^+^ cells at 2 dpi when preOLs were lineage-traced from P14, no ramified cells were present at 6 dpi (Figures 5E-G). These results suggest that a small subset of preOLs can remain at an immature stage for an extended period early postnatally.

**Figure 5.**
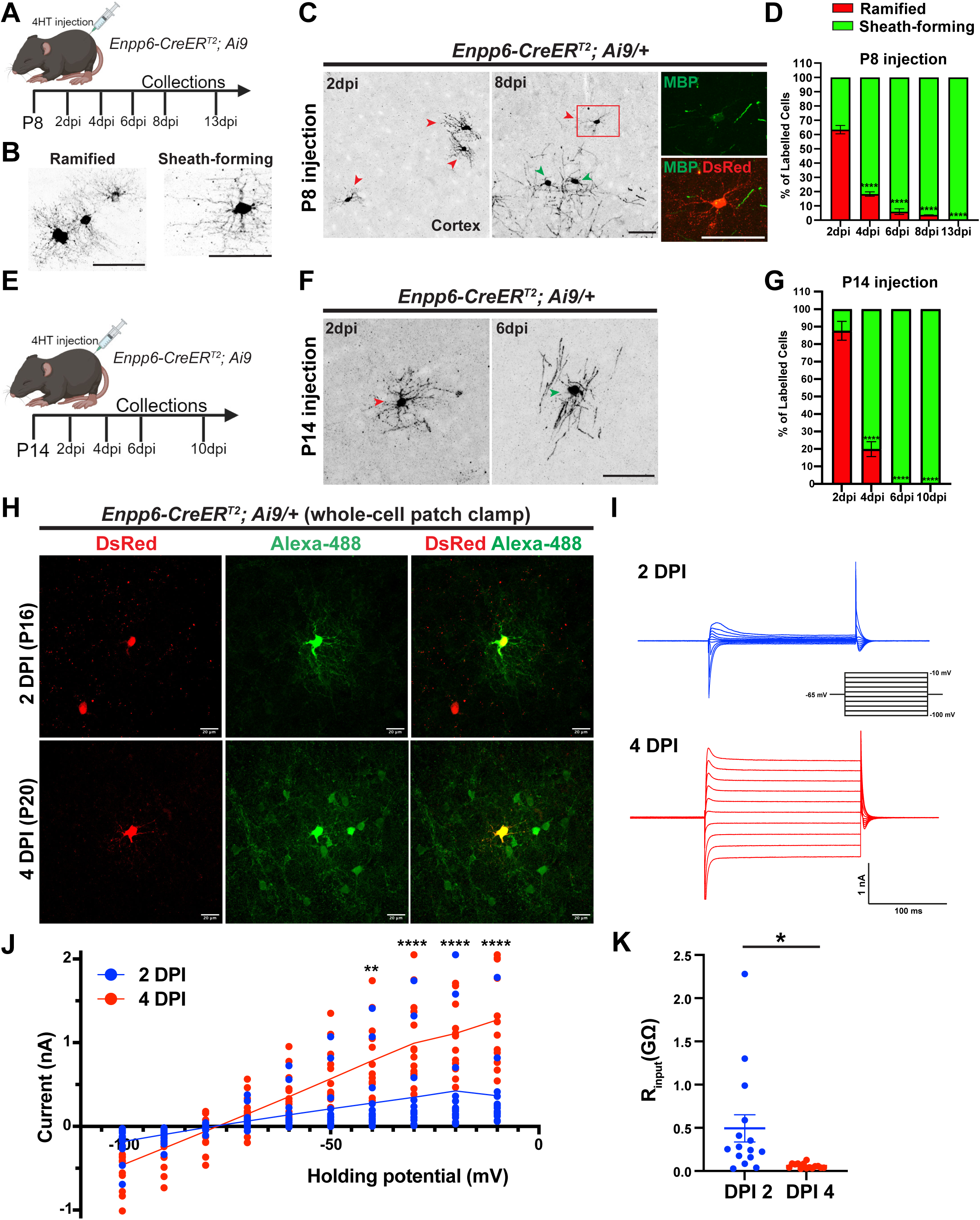
Genetically labeled preOLs exhibit distinct electrophysiological properties and undergo prolonged maturation before committing to myelination. **(A)** Schematic diagram showing the lineage tracing and characterization of preOLs in early development (P8). **(B)** Representative images of lineage-traced preOLs from P8 that undergo maturation and morphological changes. Some exhibit a “ramified” morphology (left), while others display the “sheath-forming” morphology (right). **(C)** Representative images of DsRed^+^ cells at 2 dpi and 8 dpi that were lineage-traced from P8. Red arrowheads indicate “ramified” oligodendrocytes, while green arrowheads indicate “sheath-forming” oligodendrocytes. Red inset highlights a ramified oligodendrocyte at 8 dpi (red) that exhibits low MBP expression (green). **(D)** Quantification of DsRed^+^ “ramified” and “sheath-forming” cell percentage across the cortex, lineage-traced from P8. n=3 for each timepoint. All samples were compared to the 2 dpi timepoint. **(E)** Schematic diagram showing the lineage tracing and characterization of preOLs later in development (P14). **(F)** Representative images of lineage-traced DsRed^+^ cells from P14. Red arrowhead indicates a “ramified” oligodendrocyte, while green arrowhead indicates a “sheath-forming” oligodendrocyte. **(G)** Quantification of DsRed^+^ “ramified” and “sheath-forming” cell percentage across the cortex, lineage-traced from P14. n=3 for each timepoint. All samples were compared to the 2 dpi timepoint. **(H)** DsRed^+^ cells were filled with Alexa 488 during whole-cell patch clamp recording at 2 dpi (P16; top row) and 4 dpi (P20; bottom row). **(I)** Representative traces of current responses to voltage steps from −100 mV to −10 mV (with a holding potential at - 65mV) from DsRed^+^ cells at 2 dpi (top) and 4 dpi (bottom). **(J)** Current-voltage relation (I/V) curves from 2 dpi DsRed^+^ cells (blue) and 4 dpi DsRed^+^ cells (red). **(K)** Input resistance (R_input_ =V_input_/I_output_) comparison between 2 dpi DsRed^+^ cells (blue) and 4 dpi DsRed^+^ cells. Error bars represent SEM. Each dot in (J) and (K) represents a recording from a single DsRed^+^ cell. One-way ANOVA followed by Tukey’s multiple comparisons test for (D), (G), and (J). Two-tailed *t*-test for (K). *p<0.05, **p<0.01, ***p<0.001, ****p<0.0001. Scale bars: 50 µm in (B), (C), and (F); and 20 µm in (H).

To characterize the functional maturation of preOLs, we performed whole-cell patch-clamp recordings on brain slices from *Enpp6-CreER^T2^; Ai9/+* animals at 2 and 4 dpi. Morphological visualization was achieved using Alexa Fluor 488 hydrazide in pipette solution during whole-cell recordings to label DsRed^+^ cells (Figure 5H). The amplitude of current responses to voltage steps was significantly smaller in DsRed^+^ cells at 2 dpi compared to those at 4 dpi (Figure 5I; quantified in Figure 5J). A marked decrease in the input resistance (R_input_) was observed in DsRed^+^ cells at 4 dpi relative to those at 2 dpi (Figure 5K; two-tailed *t*-test, *p*=0.0126). These findings align with previously documented electrophysiological properties characteristic of distinct OL maturation stages, including the preOL and myelinating OL stages.^40–42^ Thus, preOLs undergo developmental stage-dependent morphological and functional maturation.

### Neuronal activity acts in a critical time window to promote preOL survival and integration

Previous studies showed that neuronal activity profoundly affects oligodendrogenesis, primarily by stimulating OPC proliferation and differentiation.^13,24,27^ PreOLs have been long thought to be directly impacted by neuronal activity,^31^ as they are abundant during development and are capable of surveying and ensheathing axons, as well as undergo significant apoptosis and pruning.^43^ We hypothesized that the effect of neuronal activity would affect both OPCs and preOLs, influencing proliferation and differentiation in the former, and survival and integration in the latter (Figure 6A). To begin dissecting the roles of neuronal activity in distinct phases of oligodendrogenesis, we leveraged the preOL driver line in combination with a simple yet robust paradigm for manipulating neuronal activity.^32^ As a proof-of-principle experiment, P5 *Enpp6-CreER^T2^; Ai9/+* animals were subjected to sensory deprivation by trimming the whiskers on only one side of the mouse, leaving the other side intact as an internal control. These animals received 3 injections of 4HT during whisker trimming to label and fate-map preOLs (Figure 6B). Since whisker trimming occurred before preOL lineage tracing, the genetically labeled preOLs reflect the effects of neuronal activity on both the preOLs and the OPCs that gave rise to them. Indeed, the somatosensory cortices corresponding to the trimmed whiskers showed a significant reduction in DsRed^+^ cell density compared to the spared region (Figures 6C and 6D; quantified in Figure 6E). Importantly, no change in DsRed^+^ cell densities were observed in other brain regions, such as motor and visual cortices (Figure 6E). We also observed a significant reduction in the proportion of ramified cells over total DsRed^+^ cells in the “trimmed” somatosensory cortex, suggesting that OPC-to-mOL maturation is perturbed in the absence of sensory input (Figure 6F).

**Figure 6.**
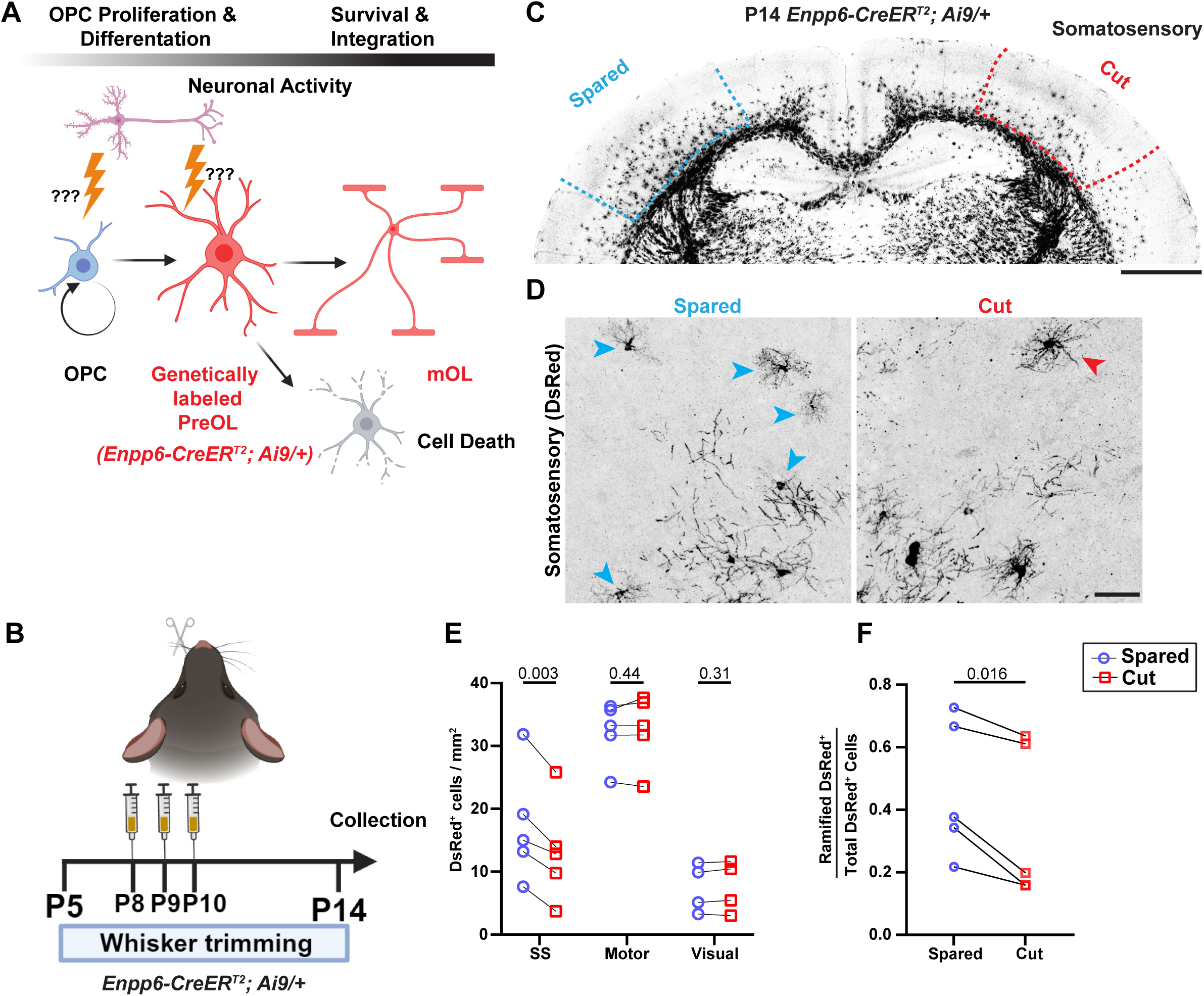
The *Enpp6-CreER^T2^* driver enables preOL fate-mapping upon perturbation of neuronal activity. **(A)** Schematic showing the potential effect of neuronal activity on oligodendrogenesis, together through its impact on oligodendrocyte progenitor cells (OPCs) as well as premyelinating oligodendrocytes (preOLs). This activity-induced change can be visualized in *Enpp6-CreER^T2^; Ai9/+* mice through genetically labeling and lineage tracing preOLs during the entire period when neural activity is perturbed. **(B)** The experimental paradigm wherein postnatal day 5 (P5) pups of *Enpp6-CreER^T2^; Ai9/+* are sensory deprived by trimming whisker of the animals on one side while leaving the whiskers on the other side intact. 100 mg/kg 4HT injections are administered to the pups on P8, P9 and P10 to label preOLs, while trimming of whisker on one side continues until the time of collection (P14). **(C)** Representative image of *Enpp6-CreER^T2^; Ai9/+* coronal section at P14 under the paradigm shown in (B). Somatosensory regions of both hemispheres, corresponding to the spared and cut whiskers, are marked with dashed lines. **(D)** High-magnification images of somatosensory cortices under the paradigm in (B) show ramified and sheath-forming DsRed^+^ cells. **(E)** Quantification of DsRed^+^ cell density in the somatosensory cortex, motor cortex, and visual cortex. n = 5 for somatosensory cortex and motor cortex. n=4 for visual cortex. **(F)** Quantification of ramified DsRed^+^ cell proportion among the total DsRed^+^ cells in somatosensory cortex following whisker trimming, based upon the paradigm shown in (B). Each dot in (E) and (F) represents a single animal. Two-tailed paired *t*-test for (E) and (F), comparing the “spared” and “cut” regions of the same animal. Scale bars: 1 mm in (C); and 50 µm in (D). SS: Somatosensory cortex; Motor: Motor Cortex; Visual: Visual Cortex.

To evaluate whether neuronal activity could be directly sensed by preOLs and regulate their survival and integration (Figure 7A), we subjected P8 *Enpp6-CreER^T2^; Ai9/+* mice to a one-sided whisker trimming paradigm with simultaneous 4HT administration, to label preOLs formed at the same time (Figure 7B). We observed a significant reduction in DsRed^+^ cell density, at P14, in the deprived somatosensory cortex as compared to the spared cortex, while no change was observed in the motor or visual cortices (Figures 7C, S6A, S6B; quantified in Figure 7D). Consistently, the density of *Mog^+^* mature oligodendrocytes, encompassing the ones formed from genetically labeled preOLs, showed a significant reduction in the deprived somatosensory cortex at P14 (Figures S6C and S6D; quantified in Figure 7F). However, both OPC numbers and MBP density in the somatosensory cortices remained unchanged (Figures S6E-F’), suggesting that reduced number of oligodendrocytes may still compensate to form the same amount of myelin during sensory deprivation.

**Figure 7.**
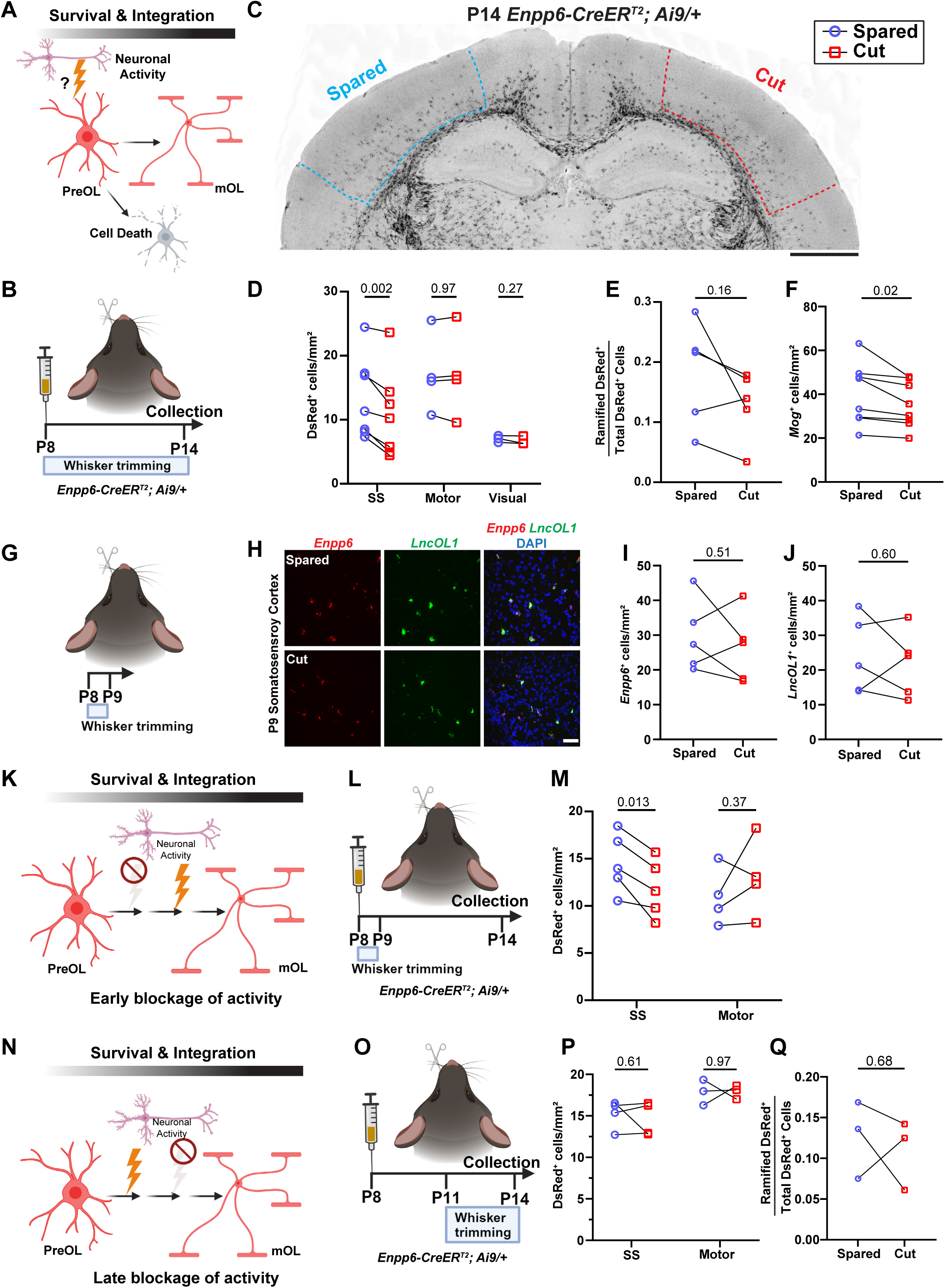
Neuronal activity acts in a critical time window to promote the survival and integration of genetically labeled premyelinating oligodendrocytes. **(A**) Schematic suggesting potential effects of neuronal activity on preOL survival and successful integration. **(B)** The experimental paradigm wherein postnatal day 8 (P8) *Enpp6-CreER^T2^; Ai9/+* pups are sensory deprived by trimming whisker of the animals on one side while leaving the whiskers on the other side intact. 150 mg/kg 4HT injection is administered to the P8 pups simultaneously with the start of whisker trimming. The whisker trimming on one side continues until the day of collection (P14). **(C)** Representative image of *Enpp6-CreER^T2^; Ai9/+* coronal section at P14 under the paradigm shown in (B). Somatosensory regions of both hemispheres, corresponding to the spared and cut whiskers, are marked with dashed lines. **(D)** Quantification of DsRed^+^ cell density in the somatosensory cortex, motor cortex, and visual cortex under the paradigm shown in (B). n=6 for somatosensory cortex, n=4 for motor cortex, and n=3 for visual cortex. **(E and F)** Quantification of ramified DsRed^+^ cell proportion among the total DsRed^+^ cells (E) and quantification of *Mog^+^* cell density (F) in the somatosensory cortex, based upon the paradigm shown in (B). **(G)** The experimental paradigm wherein C57BL/6 P8 pups are sensory deprived by trimming their whiskers only on one side while leaving the whiskers on the other side intact. The pups are collected 1 day post trimming at P9. **(H)** Representative confocal micrographs of P9 somatosensory brain area under the paradigm of (G), showing *Enpp6^+^* (red) and *LncOL1^+^*cells (green). **(I and J)** Quantification of *Enpp6*^+^ (I) and *LncOL1*^+^ (J) cell density under the paradigm of (G). n=5. **(K and L)** Schematic (K) and experimental paradigm (L) testing the effects of neuronal activity at the early phase during preOL survival and integration. *Enpp6-CreER^T2^; Ai9/+* P8 pups are sensory deprived acutely by trimming their whiskers only once on one side while leaving the whiskers on the other side intact. Meanwhile, 150 mg/kg 4HT injection is administered to P8 pups at the time of whisker trimming. **(M)** Quantification of DsRed^+^ cell density in the somatosensory and motor cortices under the paradigm of (L). n= 5. **(N and O)** Schematic (N) and experimental paradigm (O) testing the effects of neuronal activity during the late stages of preOL integration. P8 *Enpp6-CreER^T2^; Ai9/+* pups are administered with 150 mg/kg 4HT and then sensory deprived from P11 to P14 by continuously trimming the whiskers on one side while keeping the other side intact. **(P)** Quantification of DsRed^+^ cell density in the somatosensory and motor cortices under the paradigm of (N). n= 4 for somatosensory cortex and n=3 for motor cortex. **(Q)** Quantification of ramified DsRed^+^ cell proportion among the total DsRed^+^ cells in the somatosensory cortex, based upon the paradigm of (N). n= 3. Each dot in (D), (E), (F), (I), (J), (M), (P), and (Q) represents a single animal. Two-tailed paired *t*-test for (D), (E), (F), (I), (J), (M), (P), and (Q), comparing the spared and cut regions of the same animal. Scale bar: 1 mm in (C); and 50 µm in (H). SS: Somatosensory cortex; Motor: Motor Cortex; Visual: Visual Cortex.

To test whether sensory deprivation can immediately reduce preOL number, we performed whisker trimming on P8 wild-type animals and analyze *Enpp6^+^* and *LncOL1^+^* preOLs at P9 (Figure 7G). Both *Enpp6^+^* and *LncOL1^+^* preOL numbers remained unchanged after one day of sensory deprivation (Figure 7H; quantified in Figures 7I and 7J). Furthermore, we found no change in DsRed^+^ cell number in the deprived somatosensory cortex compared to the control when preOLs were lineage-traced from P8 to P11 (Figures S6G-I). Thus, sensory deprivation does not immediately alter preOL number.

As preOLs undergo prolonged maturation during early brain development (Figure 5), neuronal activity could influence either their survival at an early phase or their integration at a later stage. To determine whether neuronal activity operates in such a critical period (Figure 7K), we acutely blocked sensory input by performing a one-time whisker trimming on P8 *Enpp6-CreER^T2^; Ai9/+* mice. The whiskers were then allowed to grow back until the time of collection at P14 (Figure 7L). The deprived somatosensory cortex again showed a significant reduction in DsRed^+^ cell density compared to the spared cortex (Figure 7M). In stark contrast, when whiskers were trimmed from P11 to P14 while preOLs were fate-mapped starting from P8 (Figures 7N and 7O), we observed no change in the density of DsRed^+^ cells or in the proportion of ramified cell between the spared and deprived somatosensory cortices (Figures 7P and 7Q). Therefore, neuronal activity directly affects preOLs within a narrow time window of their early maturation, promoting their survival and successful integration.

## DISCUSSION

In this study, we generated a CreER^T2^ driver line to label subsets of preOLs throughout the CNS. Genetically labeled preOLs are postmitotic cells and exhibit distinct morphology, transcriptome, and electrophysiological properties. They emerge spatiotemporally across diverse brain regions, and sense neuronal activity in a narrow time window to survive and integrate into the local circuits. In mammals, neuronal activity-mediated changes in oligodendrogenesis have been primarily focused on OPCs, as they are known to make direct synapses with neurons.^44,45^ OPCs are evenly distributed throughout the brain, but myelination occurs in specific brain areas and neuronal circuits.^22^ Therefore, the decision about where and when to myelinate is largely determined by the bottleneck in oligodendrogenesis: the preOLs. Thus, the newly developed preOL-specific *Enpp6-CreER^T2^* mouse line fills a critical gap in the myelin biology toolkit.

PreOLs were initially defined by using an antibody raised against DM-20, an alternative spliced isoform of Plp, as immature and ramified oligodendrocytes.^4^ Due to the lack of genetic tools, they are described by various terms, such as immature oligodendrocytes, newly formed oligodendrocytes, and committed oligodendrocyte progenitors. The absence of a clear nomenclature leads to ambiguity in the precise characterization of these cells. Over the past few decades, several molecular markers have been identified to partially label preOLs. However, these RNA markers can only label preOL nuclei or cell bodies without delineating the complete morphology (e.g., *Enpp6* and *LncOL1*).^13–15^ Other markers, such as BCAS1 and Tcf7l2, are also weakly expressed by other oligodendrocyte lineage cells, including OPCs and myelinating oligodendrocytes.^12,46^ Our study shows that genetically labeled preOLs in *Enpp6-CreER^T2^; Ai9/+* mice are post-mitotic oligodendrocytes formed as OPCs exit cell cycle, encompassing both COPs and NFOLs. The majority of them exhibit a ramified morphology, while some have not yet begun expressing CC1 or MBP, two commonly used postmitotic oligodendrocyte markers (Figure 2). Our data corroborate a previous study describing *LncOL1*^+^ MBP^-^ immature preOLs.^14^ Intriguingly, our snRNA-seq analysis of genetically labeled preOLs identified a cluster termed COP1, which exhibits a unique gene expression profile likely representing the earliest stage of OPC differentiation (Figure 3).

*Enpp6* is known to be highly expressed by preOLs (*Enpp6^high^*) and lowly expressed by myelinating OLs.^13^ However, our *Enpp6-IRES-CreER^T2^* driver line predominantly labels preOLs at any time point following 4HT induction. We reason that it is likely due to the knock-in strategy, wherein the *IRES-CreER^T2^*was inserted into the 3’UTR rather than the gene coding region. This resulted in an overall reduced *CreER^T2^* expression, which is minimally detectable in myelinating OLs. Indeed, the *Enpp6-IRES-CreER^T2^*labels 10-20% of total preOLs during development. Despite reduced efficiency, this line reliably labels subsets of preOLs, as evidenced by their morphology, marker expression, transcriptomic and electrophysiological features. Moreover, this line does not preferentially label a specific subset of preOLs, because 1) labeled preOLs exhibit a “salt-and-pepper” pattern compared to the total preOL population (Figure 4), and 2) they show similar transcriptomic features to those in a published dataset encompassing preOLs (Figure 3).

PreOLs are long considered as an intermediate cell stage in oligodendrogenesis, largely based on their transient appearance and disappearance observed in both *in vitro* culture and *in vivo* long-term imaging.^3,5–7^ Our work shows that subsets of genetically labeled preOLs can exist for up to 8 days in the mouse cortex during development, challenging the stereotype of a “transient” cell stage (Figures 5 and S5). In contrast, during *in vitro* differentiation, oligodendrocytes can remain at the “preOL-like” stage for no more than 4 days.^6^ It is plausible that the presence of inhibitory cues and/or the lack of maturation cues halt preOLs in their immature state during development. This mechanism might also be relevant to demyelinating diseases, where preOLs are often in abundance but they fail to mature near lesion sites.^10^ Furthermore, the presence of immature preOLs for an extended period raises the possibility of myelination-independent functions of preOLs such as phagocytosis, which has been observed in OPCs.^47,48^ Indeed, preOLs express highly levels of the TAM receptor, Tyro3, which is critical for phagocytosis.^36,49^ It will be interesting to begin tackling such questions with this new tool for preOLs.

In humans, the brainstem, cerebellum, somatosensory and motor cortex are among the first brain regions to receive myelination.^50^ It is long hypothesized that active neuronal circuits and the ones that are critical for animals’ survival are myelinated first.^51^ In the present study, we employ the *Enpp6-CreER^T2^* line to directly visualize postmitotic oligodendrocytes that have just exited the cell cycle and to lineage-trace them, thereby mapping out oligodendrogenesis and brain-region-specific myelination during early brain development (Figure 4). Similar to humans, preOLs first appear in the mouse brainstem and cerebellum, and they quickly emerge in a posterior-to-anterior fashion.^52,53^ In addition, both instantaneous visualization and lineage tracing of preOLs show that the hippocampus, which exhibits prolonged circuitry maturation and late synapse development compared to other brain areas, is among the last brain regions to receive myelination.^54^ This data correlates well with recent findings showing that newly generated myelin reduces the synaptic plasticity of maturing circuits, thereby stabilizing the functionally important ones.^55^

It is well established that neuronal activity can impact myelination in several ways, including OPC proliferation and subsequent differentiation,^3,13,27^ the onset of axon ensheathment,^26,43^ the selection of axon for myelination,^56^ as well as the successful integration of newly generated oligodendrocytes into neuronal circuits.^23^ We provide direct evidence that neuronal activity can act on existing preOLs to promote their survival and integration. Furthermore, decreased preOL incorporation in the absence of sensory input happens early in a critical window during preOL maturation (Figure 7). This is corroborated by our single nuclei sequencing data, which shows that the “cell death” biological process is primarily enriched early in preOL maturation, specifically the COP2 cluster of preOLs (Figure 3). Hence, we speculate that preOL survival is controlled by both an intrinsic program orchestrated by the TFEB-PUMA axis,^6^ as well as by an extrinsic process in which external stimuli, such as neuronal activity, may regulate their survival.

A recent study aimed to create a preOL visualization mouse line using the *LncOL1* loci but was only able to visualize myelin-forming oligodendrocytes (MFOLs).^57^ Hence, the *Enpp6-CreER^T2^* is a useful tool for specific labeling and genetic manipulation of preOLs. It can be employed to begin addressing many longstanding questions in myelination, such as how axon selectivity is achieved during ensheathment, and which recognition molecules are required for CNS myelination. Moreover, *Enpp6-CreER^T2^* enables a clear distinction between observations originating from OPCs and those from preOLs. This is crucial for understanding demyelinating conditions in which preOL maturation is disrupted, such as preterm white matter injury and multiple sclerosis.

Despite its high specificity and being representative in demarcating preOLs, *Enpp6-CreER^T2^* labels only 10-20% of the total preOL population in the developing brain. In addition, we urge caution when using this line to conduct gene deletion at the preOL stage, given the relatively low expression levels of CreER^T2^ recombinase and the brief duration of the preOL stage. Lastly, the labeling efficiency of *Enpp6-CreER^T2^* decreases as the animal develops, with juvenile mice showing a marked reduction in preOL labeling. Therefore, it might not be the ideal tool for multi-photon live imaging experiments conducted in adult mice.

In conclusion, our study provides a powerful tool for labeling and lineage tracing subsets of preOLs during development, which we used to demonstrate the direct role of neuronal activity in regulating their survival and integration. Our work provides a framework for studying axon ensheathment, a critical early event during myelination, as well as other neuron-glia and glia-glia interactions where preOLs might play a role.

## ACKNOWLEDGEMENTS

We thank the members of the Sun laboratory for assistance and discussions. This work was supported by the UT Southwestern Endowed Scholarship (L.O.S.), the NIH Director’s New Innovator Award (4DP2MH129988-02, L.O.S.), the Chan Zuckerberg Initiative Ben Barres Early Career Acceleration Award (L.O.S.), the Klingenstein-Simons Foundation Fellowship Award in Neuroscience (L.O.S.), the NIH grant U19NS123719 (G.Y.), and R01 DC018797 (J.H.K.). L.O.S. is a Southwestern Medical Foundation Scholar in Biomedical Research.

## AUTHOR CONTRIBUTIONS

Conceptualization, A.B., L.O.S.; Methodology, A.B., P.T., S.W., H.G.B., D.B., T.S., Z.Y., C.X., J.H.K., G.Y., L.O.S.; Investigation, A.B., P.T., S.W., H.G.B., Z.Y., C. X., J.H.K., G.Y., L.O.S.; Writing, A.B., C.X., J.H.K., G.Y., L.O.S.; Funding Acquisition, C.X., G.Y., J.H.K., L.O.S.; Supervision, C.X., J.H.K., G.Y., L.O.S.

## SUPPLEMENTAL INFORMATION

### Other Supplemental Items

**Supplementary Table 1 (Table S1): Microinjection details for *Enpp6-IRES-CreER^T2^* mice generation.**

The table lists the total number of embryos injected in two rounds of injections, the number of viable embryos implanted, the number of live pups born, and the number of site-specific knock-in animals.

**Supplementary Table 2 (Table S2): Genes enriched in different clusters and their expression, extracted from Marques, et al. (Science, 2016) dataset, used for similarity score analysis in Figure S2, related to Figure 3.**

Each table in the excel sheet represents the cluster: OPC, COP, NFOL, MFOL and MOL. In the table, each column represents: Gene, p-value (calculated using Wilcox Rank Sum test), the average log_2_(fold change) (Fold Change is calculated comparing expression of gene in the given cluster with the expression of gene in the rest of clusters), Percentage of cells expressing the gene in the main cluster analyzed, Percentage of cells expressing the gene in the rest of the clusters and adjusted p-value (multiple testing correction performed using Bonferroni correction method).

**Supplementary Table 3 (Table S3): Differentially Enriched Genes (DEGs) for each cluster in single nuclei sequencing of preOL lineage cells, related to Figure 3.**

Each table in the excel sheet represents the clusters: COP1, COP2, NFOL1, NFOL2, NFOL3, NFOL4, MFOL, MOL, CPC1, CPC2, IRC. The enriched genes are filtered based on z-score > 2.6 with p-value < 0.01. In the table, each column represents: Gene, z-score (calculated using p-value from Wilcox Rank Sum test), the average log_2_(fold change) (Fold Change is calculated comparing expression of gene in the given cluster with the expression of gene in the rest of clusters), Percentage of cells expressing the gene in the main cluster analyzed and Percentage of cells expressing the gene in the rest of the clusters.

### Supplemental Figure Legends

**Figure S1, related to Figure 1.**
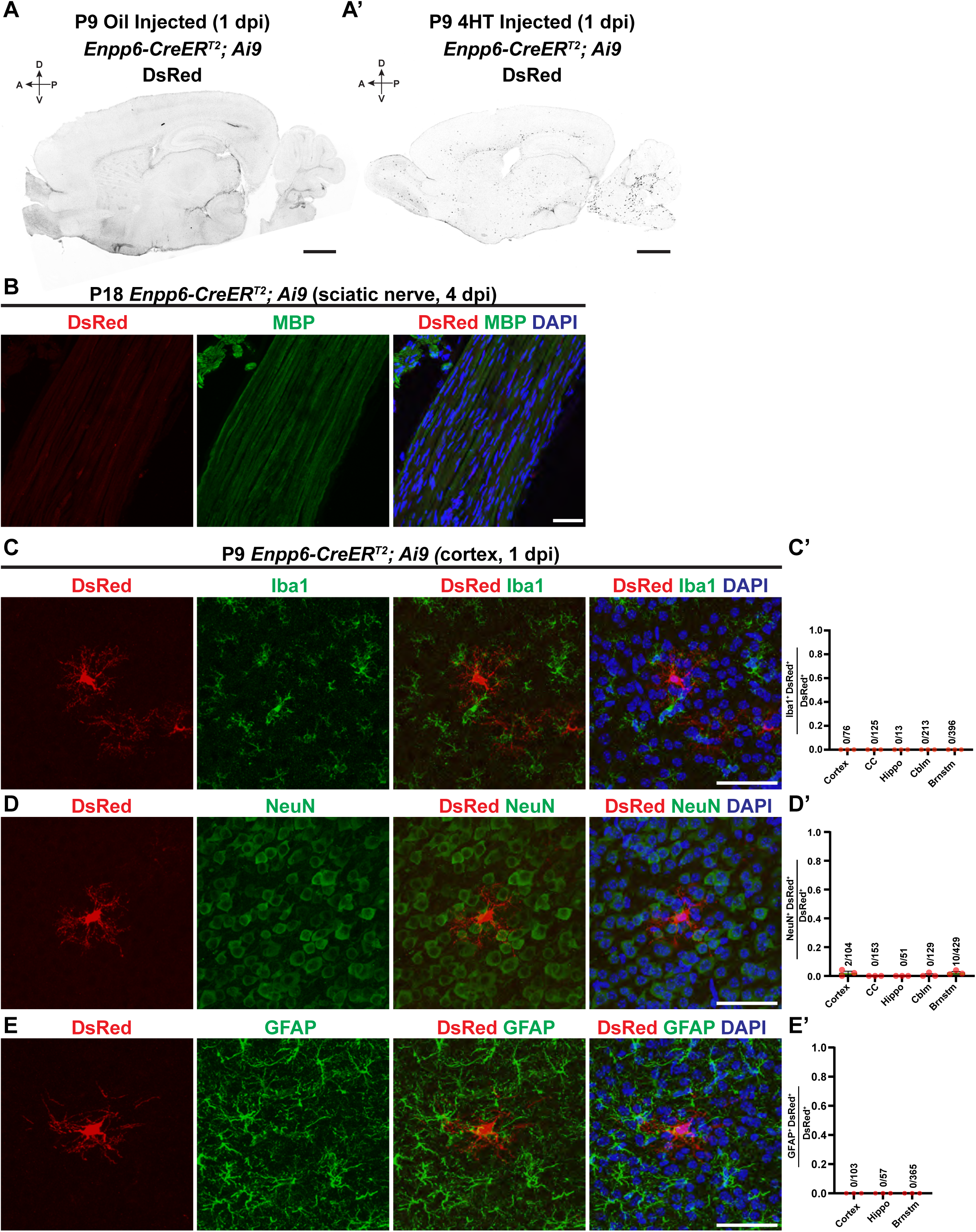
*Enpp6-IRES-CreER^T2^* shows no leakage of expression into the peripheral nervous system, astrocytes, or microglia in the brain, and exhibits minimal CreER^T2^ expression in neurons. **(A and A’)** Sagittal brain sections of postnatal day 9 (P9) *Enpp6-CreER^T2^; Ai9/+* mice, 1 day post injection (dpi) with sunflower oil (A) or 150 mg/kg 4-hydroxytamoxifen (4HT; A’). The DsRed^+^ cells were visualized by immunohistochemistry using an anti-DsRed antibody. A, anterior; P, posterior; D, dorsal; V, ventral. **(B)** Representative images of P18 *Enpp6-CreER^T2^; Ai9/+* sciatic nerve at 4 dpi of 100 mg/kg 4HT stained by DsRed (red) and MBP (green) antibodies. **(C and C’)** Representative images (C) and quantification (C’) of Iba1^+^ DsRed^+^ cell proportion among the total DsRed^+^ cells in P9 *Enpp6-CreER^T2^; Ai9/+* cortices at 1 dpi. **(D and D’)** Representative images (D) and quantification (D’) of NeuN^+^ DsRed^+^ cell proportion among the total DsRed^+^ cells in P9 *Enpp6-CreER^T2^; Ai9/+* cortices at 1 dpi. **(E and E’)** Representative images (E) and quantification (E’) of GFAP^+^ DsRed^+^ cell proportion among the total DsRed^+^ cells in P9 *Enpp6-CreER^T2^; Ai9/+* cortices at 1 dpi. Error bars represent SEM. Scale bars: 1 mm in (A) and (A’); 50 µm in (B), (C), (D), and (E). Close circles in (C’), (D’), and (E’) represent individual animals. n=3 animals per category. CC: Corpus Callosum; Hippo: Hippocampus; Cblm: Cerebellum; Brnstm: Brainstem.

**Figure S2, related to Figure 3.**
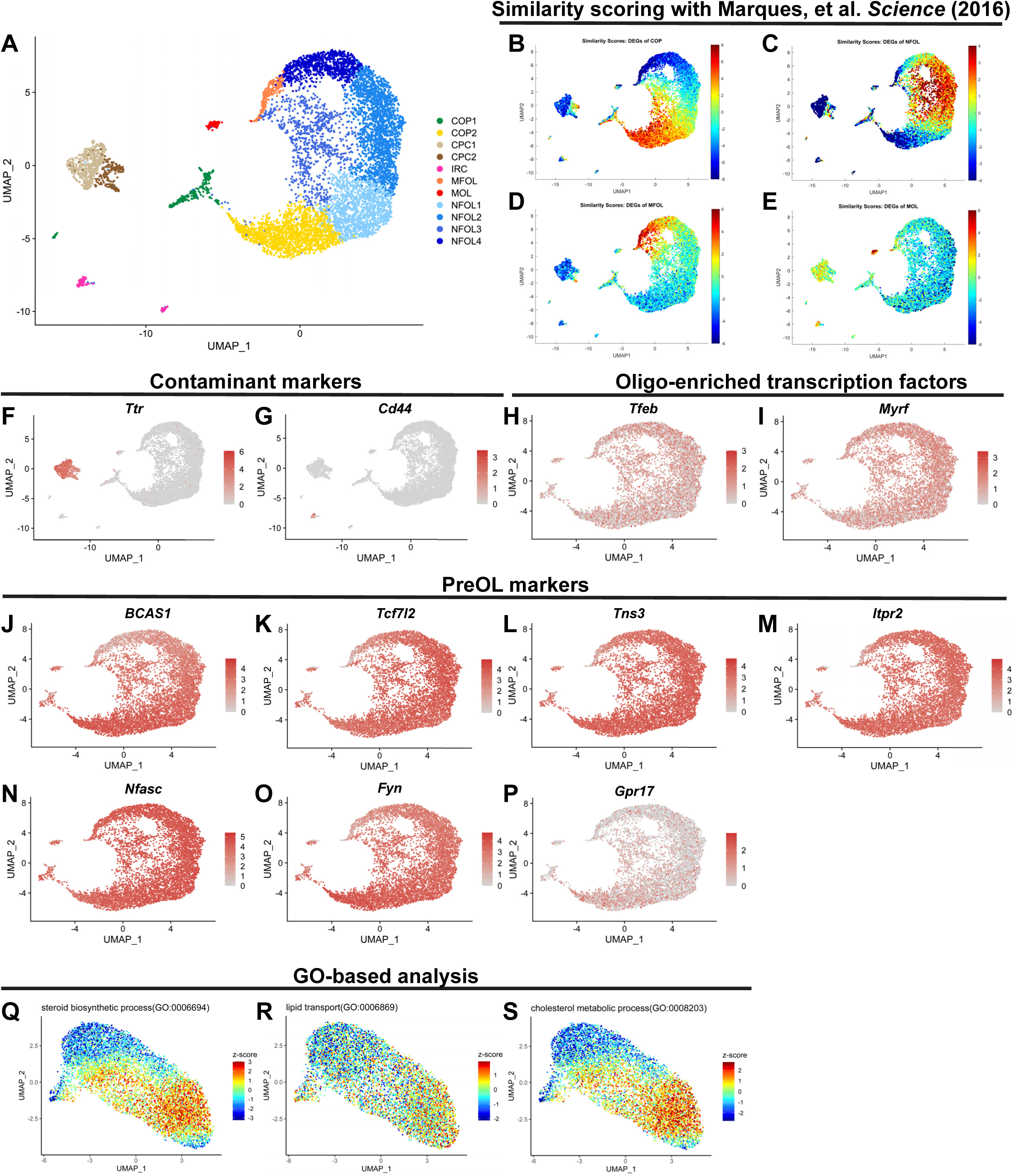
Genetically labeled preOLs express canonical preOL markers and show differentiation-stage-dependent expression of lipid metabolism pathways. **(A)** Uniform Manifold Approximation and Projection (UMAP) plots showing the preOL lineage composition, based on Seurat-based clustering of GFP^+^ cells from P9 *Enpp6-CreER^T2^; Sun1-GFP/+* mouse brains at 1 dpi. **(B-E)** Comparisons between clusters shown in (A) and a publicly available dataset from Marques, et al. *Science* (2016) using similarity scores calculated by TySim package. Clusters of COP (B), NFOL (C), MFOL (D), and MOL (E) from Marques, et al., *Science* (2016) are compared to the ones from genetically labeled preOLs. See details in Methods and Materials regarding how similarity scores are calculated and compared between these two datasets. **(F and G)** UMAP showing the expression of *Ttr* (F), a choroid plexus marker, and the expression of *Cd44* (G), a blood cell marker, in several contaminant clusters. **(H and I)** UMAP showing the expression of oligo-enriched transcription factors, *Tfeb* (H) and *Myrf* (I), across genetically labeled preOL lineage. **(J-P)** UMAP showing the expression of previously known preOL markers, such as *BCAS1* (J), *Tcf7l2* (K). *Tns3* (L), *Itpr2* (M), *Nfasc* (N), *Fyn* (O), and *Gpr17* (P), in genetically labeled preOLs. **(Q-S)** UMAP showing GO-term-based clustering of genetically labeled preOLs with a focus on lipid metabolism, based upon the annotation shown in Figure 3B. The UMAPs show z-scores of “steroid biosynthesis process” (Q), “lipid transport” (R), and “cholesterol metabolic process” (S) in preOL lineage cells.

**Figure S3, related to Figure 3.**
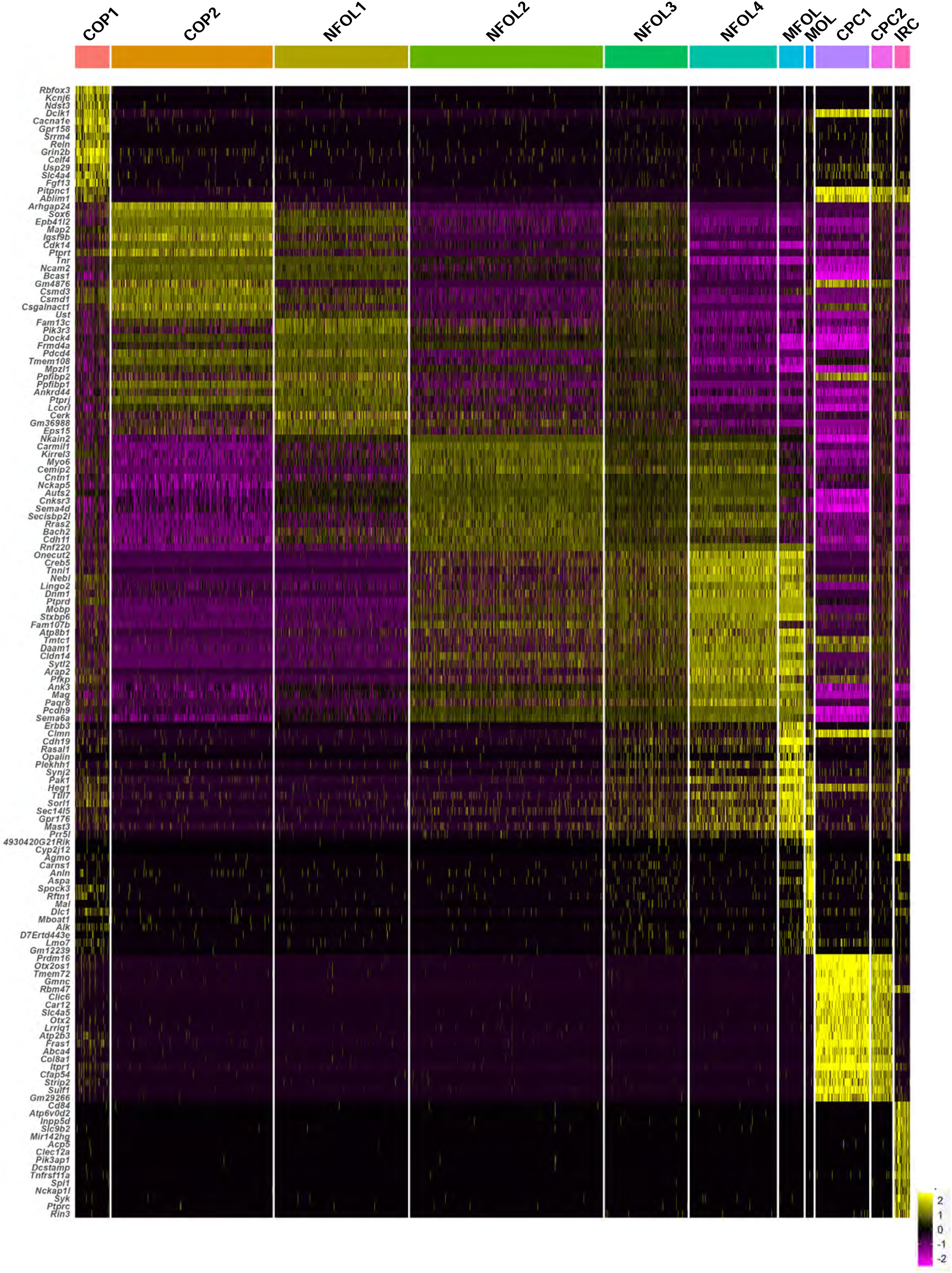
Heatmap showing the expression of cluster-enriched genes across multiple clusters of genetically labeled preOLs. Top 15 cluster-enriched genes are obtained through Seurat package analysis. The heatmap is plotted with columns representing cells of each cluster in genetically labeled preOL lineage from Figures 3 and S2, including COP1, COP2, NFOL1, NFOL2, NFOL3, NFOL4, MFOL, MOL, CPC1, CPC2, and IRC. Each row represents a gene expressed in the clusters. The legend indicates gene expression levels, with yellow representing high expression and magenta representing low expression.

**Figure S4, related to Figure 4.**
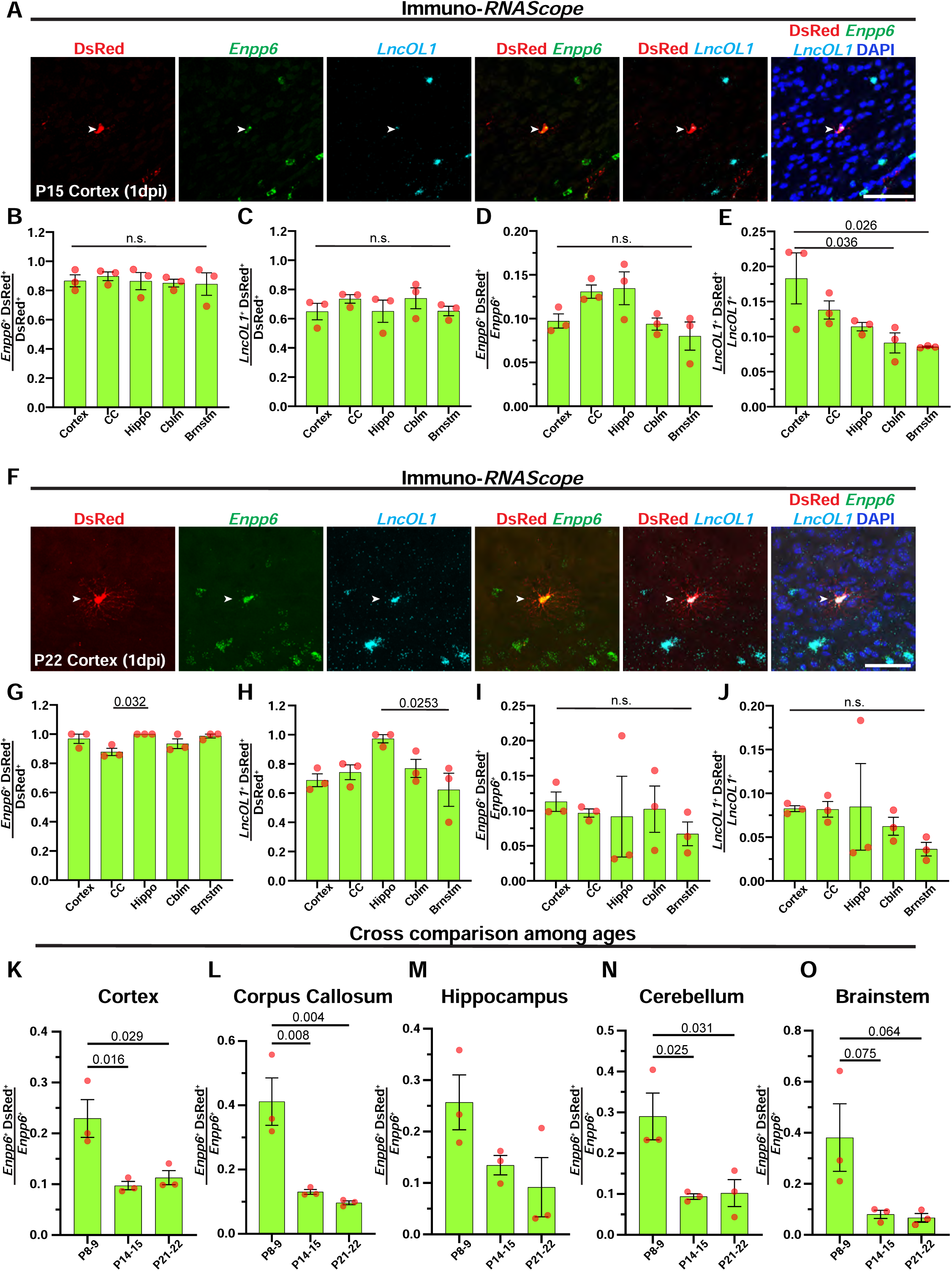
*Enpp6-IRES-CreER^T2^* labels subsets of preOLs across diverse brain regions, with the highest labeling efficiency observed during early development. **(A)** Representative images of immunohistochemistry combined with RNAScope (immuno-*RNAScope*) using DsRed antibody (red), *Enpp6 in situ* probe (green), and *LncOL1 in situ* probe (cyan) in P15 *Enpp6-CreER^T2^; Ai9/+* cortex at 1 dpi. The arrowheads indicate a DsRed^+^ cell co-expressing *Enpp6* and *LncOL1*. **(B and C)** Quantification of *Enpp6*^+^ DsRed^+^ (B) and *LncOL1*^+^ DsRed^+^ (C) cell proportions among the total DsRed^+^ cells in P15 *Enpp6-CreER^T2^; Ai9/+* cortices at 1 dpi. **(D and E)** Quantification of *Enpp6*^+^ DsRed^+^ (D) and *LncOL1*^+^ DsRed^+^ (E) cell proportions among the total *Enpp6*^+^ cells and *LncOL1*^+^ cells, respectively, in P15 *Enpp6-CreER^T2^; Ai9/+* cortices at 1 dpi. **(F)** Representative images of immuno-*RNAscope* using DsRed antibody (red), *Enpp6 in situ* probe (green), and *LncOL1 in situ* probe (cyan) in P22 *Enpp6-CreER^T2^; Ai9/+* cortex at 1 dpi. The arrowheads indicate a DsRed^+^ cell co-expressing *Enpp6* and *LncOL1*. **(G and H)** Quantification of *Enpp6*^+^ DsRed^+^ (G) and *LncOL1*^+^ DsRed^+^ (H) cell proportions among the total DsRed^+^ cells in P22 *Enpp6-CreER^T2^; Ai9/+* cortices at 1 dpi. **(I and J)** Quantification of *Enpp6*^+^ DsRed^+^ (I) and *LncOL1*^+^ DsRed^+^ (J) cell proportions among the total *Enpp6*^+^ cells and *LncOL1*^+^ cells, respectively, in P22 *Enpp6-CreER^T2^; Ai9/+* cortices at 1 dpi. **(K-O)** Comparison of *Enpp6*^+^ DsRed^+^ cell proportion among the total *Enpp6^+^* cells at multiple timepoints (P9, P14, and P22) in the cortex (K), corpus callosum (L), hippocampus (M), cerebellum (N) and brainstem (O) of *Enpp6-CreER^T2^; Ai9/+* at 1 dpi. Error bars represent SEM. One-way ANOVA followed by Tukey’s multiple comparisons test for all comparisons in (B)-(E) and (G)-(O). Close circles in (B)-(E) and (G)-(O) represent individual animals. n=3 animals per category. Scale bars: 50 µm in (A) and (F). CC: Corpus Callosum; Hippo: Hippocampus; Cblm: Cerebellum; Brnstm: Brainstem.

**Figure S5, related to Figure 5.**
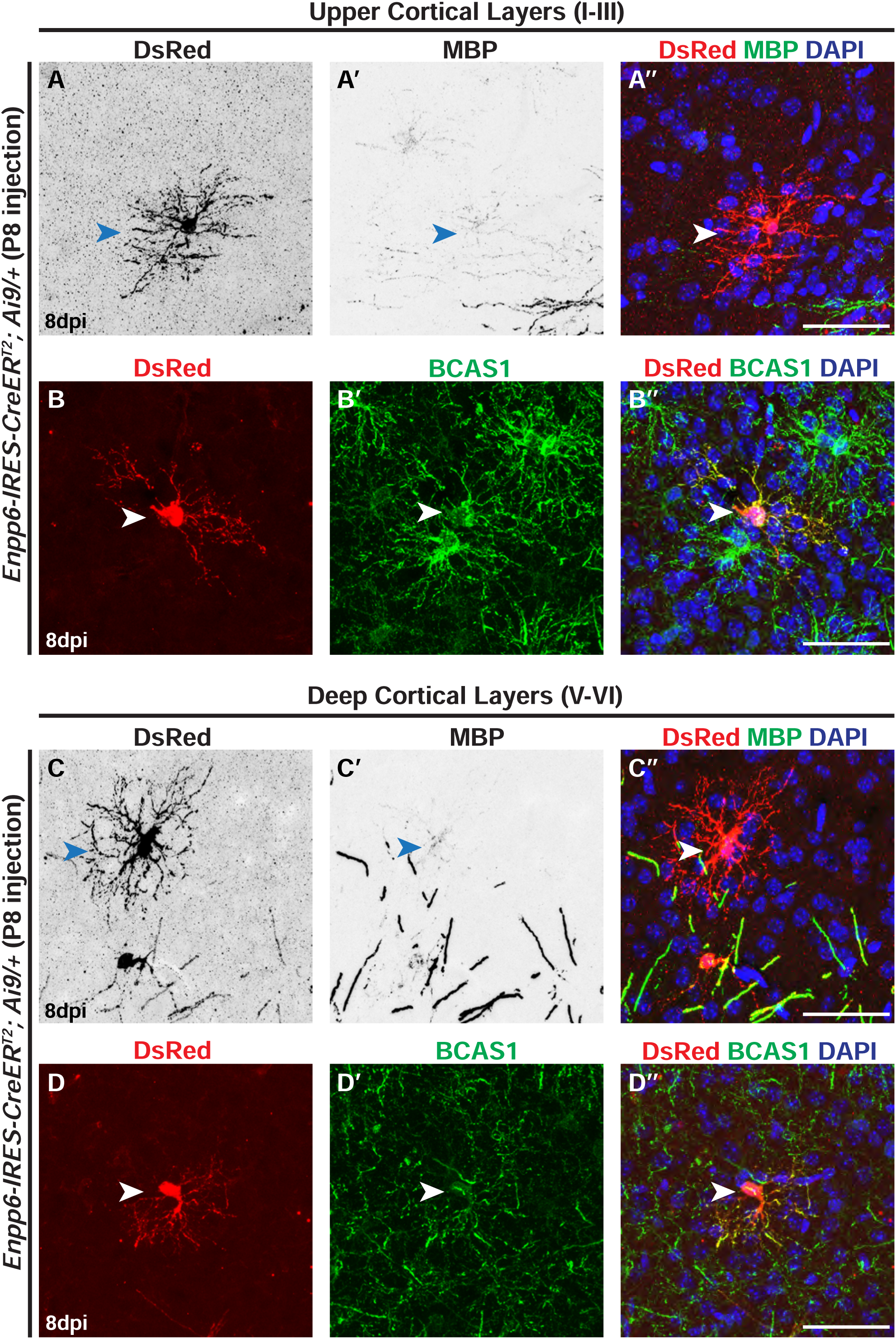
Subsets of genetically labeled preOLs exhibit a prolonged, ramified morphology during early development. **(A-A”)** Representative images of a genetically labeled preOL (A) in the upper cortical layer of a P16 *Enpp6-CreER^T2^; Ai9/+* mouse at 8 dpi, showing its low-level MBP expression (A’ and A”). **(B-B”)** Representative images of a genetically labeled preOL (B) in the upper cortical layer of a P16 *Enpp6-CreER^T2^; Ai9/+* mouse at 8 dpi, showing co-expression of BCAS1 (B’ and B”). **(C-C”)** Representative images of a genetically labeled preOL (C) in the deep cortical layer of a P16 *Enpp6-CreER^T2^; Ai9/+* mouse at 8 dpi, showing its low-level MBP expression (C’ and C”). **(D-D”)** Representative images of a genetically labeled preOL (D) in the deep cortical layer of a P16 *Enpp6-CreER^T2^; Ai9/+* mouse at 8 dpi, showing co-expression of BCAS1 (D’ and D”). Scale bars: 50 µm in (A”), (B”), (C”), and (D”).

**Figure S6, related to Figure 6 and Figure 7.**
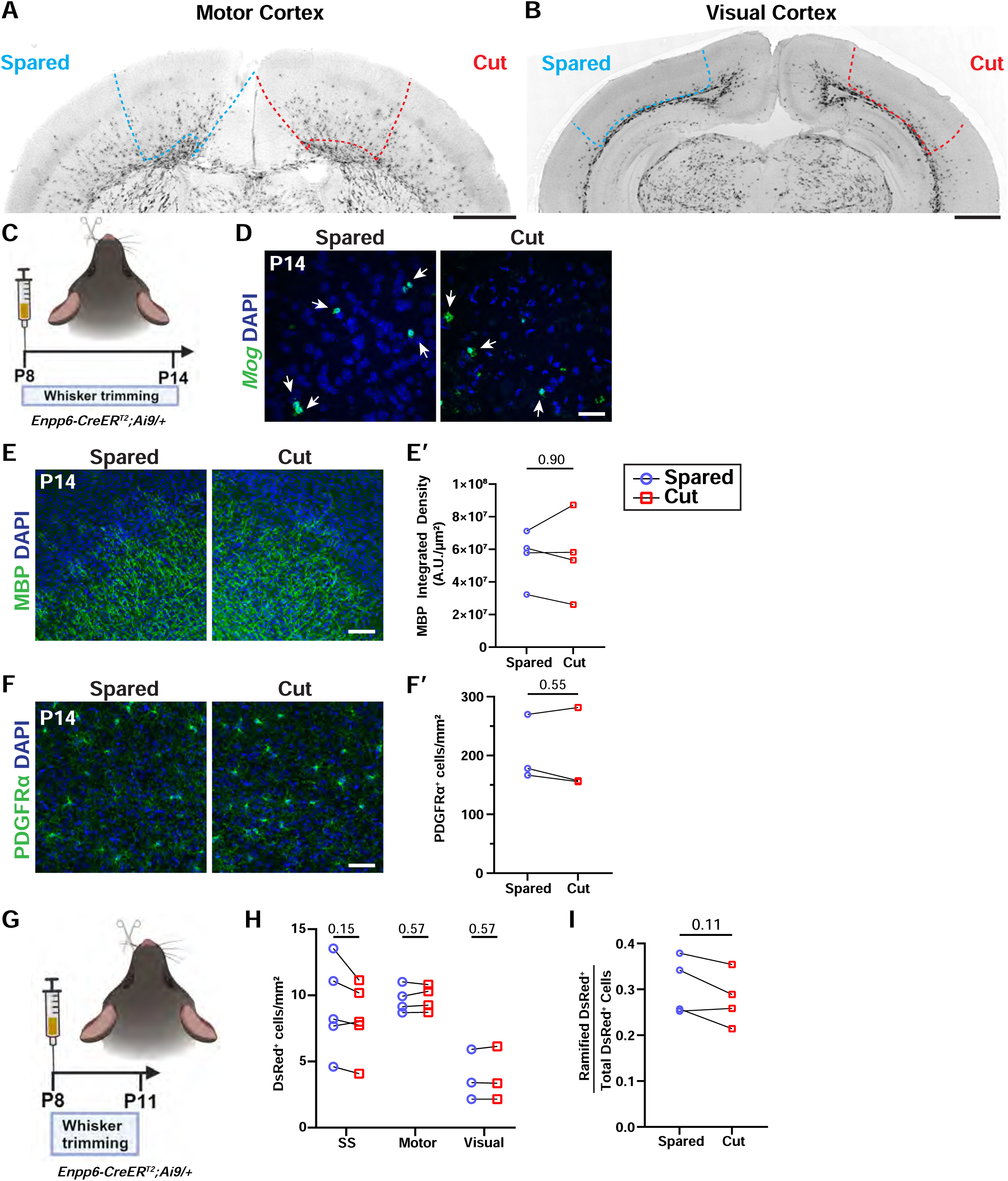
Fate-mapping of genetically labeled preOLs in multiple brain regions under a sensory deprivation paradigm. **(A and B)** Representative images of P14 *Enpp6-CreER^T2^; Ai9/+* coronal brain sections harboring the motor cortex (A) and visual cortex (B). The demarcated areas with dash lines in the motor cortex (A) and visual cortex (B) represent the brain regions used for quantifications in Figures 6E, 7D, 7M, 7P, and S6H. **(C)** The experimental paradigm wherein postnatal day 8 (P8) *Enpp6-CreER^T2^; Ai9/+* pups are sensory deprived by trimming whisker of the animals on one side while leaving the whiskers on the other side intact. 150 mg/kg 4-hydroxytamoxifen (4HT) injection is administered to the P8 pup simultaneously with the start of whisker trimming. The whisker trimming on one side continues until the day of collection (P14). **(D)** Representative images of *Mog^+^* cells in the somatosensory cortex following whisker trimming, based on the paradigm shown in (C). See quantification in Figure 7F. **(E and E’)** Representative images (E) and quantification (E’) of MBP Integrated Density in the somatosensory cortex following whisker trimming, based on the paradigm shown in (C). **(F and F’)** Representative images (F) and quantification (F’) of PDGFRα^+^ cell density in the somatosensory cortex following whisker trimming, based on the paradigm shown in (C). **(G)** The experimental paradigm wherein P8 *Enpp6-CreER^T2^; Ai9/+* pups are sensory deprived by trimming whisker of the animals on one side while leaving the whiskers on the other side intact. 150 mg/kg 4HT is administered to the P8 pup simultaneously with the start of whisker trimming. The whisker trimming on one side continues until the day of collection (P11). **(H)** Quantification of DsRed^+^ cell density in the somatosensory cortex, motor cortex, and visual cortex under the paradigm shown in (G). n=5 for somatosensory cortex, n=4 for motor cortex, and n=3 for visual cortex. **(I)** Quantification of ramified DsRed^+^ cell proportion among the total DsRed^+^ cells in the somatosensory cortex following whisker trimming, based on the paradigm shown in (G). Each circle in (E’), (F’), (H), and (I) represents a single animal. Two-tailed paired *t*-test for (E’), (F’), (H), and (I), comparing the “spared” and “cut” regions of the same animal. Scale bars: 1 mm in (A) and (B); 50 µm in (D) and (F); and 100 µm in (E). SS: Somatosensory cortex; Motor: Motor Cortex; Visual: Visual Cortex.

## MATERIALS AND METHODS

### Animals

All experiments using animals complied with the Guide for Care and Use of Laboratory Animals and protocols used were approved by Institutional Animal Care and Use Committee (IACUC) at the University of Texas Southwestern Medical Center. Mice were housed on a 12:12 light-dark cycle with food and water available *ad libitum*. The day of birth in this study was designated postnatal day 0 (P0) and all the developmental stages were specified in figures and figure legends. Both male and female animals were used for the study.

*Enpp6-IRES-CreER^T2^* transgenic line was generated using the *Easi*-CRISPR method as described previously.^58^ Briefly, sgRNAs were designed using Broad Institute sgRNA design algorithm. Pure sgRNA (Synthego) was combined with recombinant Alt-R S.p. Cas9 nuclease V3 protein (Integrated DNA Technologies, Cat#1081058) and was injected in C57BL/6J single cell embryos with ssDNA, containing *IRES-CreER^T2^* flanked by homologous regions of the *Enpp6* 3’UTR, as the repair template. After microinjections, embryos were incubated overnight in CO_2_ incubator at 37°C. Surviving embryos were transplanted in pseudo-pregnant CD1 female mice to obtain live pups. The details of injected mice and number of surviving pups are provided in Table S1. Chimeric mice were genotyped by sequencing *Enpp6* 3’UTR regions and the inserted IRES-CreER^T2^ regions. The genotyping primers are: Enpp6_Fw1 5’-GCTATGGCAGGAAGAAAGAGATA-3’, Enpp6_Fw2 5’-ACTTTGTATAGCTGGCCCTATG-3’, Enpp6_Fw3 5’-TCGATGCAACGAGTGATGAG-3’, Enpp6_Fw4 5’-GATTTACGGCGCTAAGGATGA-3’, Enpp6_Fw5 5’-GCTCTACTTCATCGCATTCCT-3’, Enpp6_Rv1 5’-AGACGGCAATATGGTGGAAA-3’, Enpp6_Rv2 5’-TCTTGCGAACCTCATCACTC-3’, Enpp6_Rv3 5’-CTTGCATGATCTCCGGTATTGA-3’, Enpp6_Rv4 5’-AGCTGTGGCAGGGAAAC-3’ and Enpp6_Rv5 5’-GGGAAGAACCAGACAACACATA-3’. Founder mice were backcrossed with wildtype C57BL/6J mice for 10 generations.

*Enpp6-IRES-CreER^T2^* were crossed with reporter mice for characterization. Reporter mouse lines procured from the Jackson Laboratory were the Ai9 reporter (stock #007909) and the *Sun1-GFP* reporter (stock #021039).

### Immunohistochemistry

Mice were euthanized with CO_2_ and immediately perfused with chilled PBS (Sigma-Aldrich, Cat#PPB006) and 4% PFA (EMS, Cat#15710), made in 1X PBS. The brains were post-fixed in a 4% PFA/PBS solution overnight at 4°C, then transferred to a 30% sucrose solution in 1X PBS at 4°C for 24 hours. Blocks were made by removing the brain tissue from sucrose and immersing them in embedding medium (Fisher Healthcare Tissue-Plus O.C.T. Compound, Cat#23-730-571) and freezing the sample on dry ice. Cryosectioning was done using a Leica CM1950 Cryostat. Sagittal or coronal sections of 18 µm thickness were used for immunostaining while 12 µm thick sections were used for immunostaining combined with *in situ* hybridization. Optic nerves and spinal cords were collected by the same perfusion methods and were processed similarly. Optic nerves were sectioned at 12 µm thickness for immunostaining while spinal cords were sectioned at 18 µm thickness for staining.

For immunostaining, the sections were incubated with either 10% normal goat serum (Jackson ImmunoResearch, Cat#005-000-121) or 10% normal donkey serum (Jackson ImmunoResearch, Cat#017-000-121) in PBS supplemented with 0.1% Triton X-100 (PBST; Sigma-Aldrich, Cat#T8787) for 30 minutes at room temperature in a humified chamber. The primary antibodies were diluted in PBST containing 2% normal goat serum or normal donkey serum. Sections were incubated in primary antibodies overnight at 4°C. The sections were washed with PBST 10 minutes for 4 times at room temperature. Sections were incubated with AlexaFluor 488- or 555-conjugated secondary antibodies (ThermoFisher, 1:1000) and DAPI (ThermoFisher, 5 μg/mL, Cat#62247), diluted in PBST containing 2% serum for 1 hour at room temperature. The sections were washed 4 times with PBST and mounted using VectaShield Hardset Antifade mounting medium (Vector Laboratories, Cat#H-1700-10). Images were taken using a Keyence BZ-810 all-in-one fluorescence microscope or a Zeiss LSM 700 inverted confocal microscope.

Primary antibodies used in this study include: Rabbit Anti-DsRed (Clonetech, 1:1000, Cat#632496), Goat Anti-DsRed (Biorybt, 1:1000, Cat#orb11618), Goat Anti-Olig2 (R&D systems, 1:1000, Cat#AF2418SP), Goat Anti-Sox10 (R&D systems, 1:1000, Cat#AF2864), Mouse Anti-CC1 (Millipore, 1:200, Cat#OP80), Rat Anti-MBP (abcam, 1:4000, Cat#7349), Rabbit Anti-NeuN (Cell Signaling, 1:1000, Cat#12943), Chicken Anti-GFAP (Aves Lab, 1:2000, Cat#GFAP, AB_2313547), Rabbit Anti-Iba1 (Wako Chemicals, 1:1000, Cat#019-19741), Rabbit Anti-BCAS1 (Synaptic Systems, 1:1000, Cat#445003) and Rabbit Anti-PDGFRα (abcam, 1:1000, Cat#ab203491).

### 4-hydroxytamoxifen preparation and injections

4-hydroxytamoxifen (4HT; Sigma-Aldrich, Cat#H7904-25mg) was dissolved in 500 µL 100% ethanol at 55°C for 5-10 minutes. After repeated vortexing to dissolve the 4HT completely, the solution was transferred to 2 ml Chen Oil (4:1, Sunflower Oil: Castor Oil). The solution was vortexed and incubated at 55°C for 5-10 minutes to completely dissolve the 4HT. Aliquots were made and stored in −80°C to be used later.

Before use, the 4HT aliquot was thawed at 55°C for 5-10 minutes for intraperitoneal injections in mice. Thawed 4HT was never left at room temperature for more than 30 minutes before injections.

For characterization of *Enpp6-CreER^T2^; Ai9/+,* pups were injected with 150 mg/kg 4HT intraperitoneally. These mice were collected 1 dpi. For lineage tracing experiments (Figures 4 and 5), pups were injected with 100 mg/kg 4HT intraperitoneally.

### EdU pulse labelling

5-ethynyl-2-deoxyuridine (EdU; Sigma-Aldrich, Cat#900584) was dissolved at 10 mg/ml in DMSO (Sigma-Aldrich, Cat#D8418). EdU pulse labeling was performed by injecting a 10 mg/ml EdU solution at a dosage of 50 mg/kg in pups. If tamoxifen injections were to be combined with EdU injections, it was injected 5-10 minutes after EdU injections to allow mice to recover. Tamoxifen (Sigma-Aldrich, Cat#T5648) used was dissolved in sunflower oil at 10 mg/ml concentration. EdU^+^ cells were revealed by the Click-iT EdU Alexa Fluor 488 imaging kit (ThermoFisher, Cat#C10337) as per manufacturer’s protocol.

### Immunohistochemistry combined with *in situ* hybridization

12 µm thick cryosections were cut from sucrose-preserved brain blocks and were processed using the RNAscope Multiplex Fluorescent reagent kit v2 with TSA Vivid dyes (ACDbio, Cat#323270). *Enpp6* probe (ACDbio, Cat#511021-C2), *9630013A20Rik* probe (*LncOL1*; ACDbio, Cat#526741-C3) and *Mog* probe (ACDbio, Cat#492981) were used as per manufacturer’s instructions. Representative images were taken using a Keyence BZ 810 all-in-one microscope and a Zeiss LSM700 inverted confocal microscope.

### Nuclei preparation and single-nuclei RNA sequencing (snRNA-seq)

To genetically label the nuclei of *Enpp6^+^* oligodendrocytes, two P8 *Enpp6-IRES-CreER^T2^*; *Sun1-GFP* pups were administered with 150 mg/kg 4HT. 1 day post injection, brains from P9 *Enpp6-IRES-CreER^T2^*; *Sun1-GFP* pups were collected as per 10X genomics protocol. Briefly, the brains from two pups were micro-dissected, diced, and incubated in lysis buffer (10 mM Tris HCl pH 7.4, 10 mM NaCl, 3 mM MgCl_2_, 0.1% NP40) for 15-20 minutes. The brain tissues were triturated and spun down at 500g at 4°C for 5 minutes. Repeated washes were performed using Nuclei Wash Buffer (1% BSA, RNase Inhibitor 0.2 U/µl, 1XPBS). Nuclei were resuspended in 1 ml Nuclei Wash Buffer and incubated with 120 µl myelin removal beads (Miltenyi Biotech, Cat#130-096-733) for 15 minutes. The nuclei were diluted with an additional 10 ml Nuclei Wash Buffer. After resuspension, nuclei were spun down at 500g at 4°C for 5 minutes. Pelleted nuclei were resuspended in 2 ml nuclei wash buffer and divided into 2 LS Columns (Miltenyi Biotech, Cat#130-042-401). Nuclei were further resuspended in nuclei wash buffer and placed on a sucrose cushion (Sigma-Aldrich, Cat#NUC-201). Nuclei were spun down at 13000g at 4°C for 45 minutes in swinging bucket centrifuge, and they were resuspended in nuclei wash buffer. Resuspended nuclei were subjected to flow cytometry to sort GFP^+^ nuclei using BD Aria II sorter. After sorting, GFP^+^ nuclei were subjected to Chromium Next GEM Single Cell Gene Expression pipeline for next-generation sequencing as per manufacturer’s instructions.

### Single nuclei sequencing analysis

The alignment of the single nuclei sequencing (snRNA-seq) raw data was performed utilizing CellRanger-ARC-2.0.0 developed by 10x Genomics. Briefly, the “count” command was employed to process the raw sequencing data, with the Mus musculus reference genome (mm10) applied in the alignment process. The reference genome was obtained from https://cf.10xgenomics.com/supp/cell-arc/refdata-cellranger-arc-mm10-2020-A-2.0.0.tar.gz. The “Seurat” package (version: 5.0.1) was used for preprocessing GEX data.^59^ Initially, gene counts were normalized to total expression and log-transformed for cells passing quality control (QC) thresholds (*Log_2__CountRNA < 17*, *Log_2__CountRNA > 10*, *nFeature_RNA > 400*, *nFeature_RNA < 6000*, *percent.mt < 5/100*). In the analysis, highly variable genes were identified, and the top 2000 among them were used as input for principal component analysis (PCA) on scaled data. The top principal components (PCs) were then selected to compute the distance metric. This metric was utilized for unsupervised cell-clustering analysis. Annotation based on typical marker genes’ expression levels was performed on the results. Lastly, the results were visualized in a low-dimensional projection using Uniform Manifold Approximation and Projection (UMAP).^60^

To identify differentially expressed genes (DEGs) within each group, DEG analysis was conducted on the snRNA-seq data using the *FindAllMarkers* function in the “Seurat” package. Wilcoxon rank sum tests were performed with parameters: *logfc.threshold = 1* and *min.pct = 0.25* for a relaxed threshold; *logfc.threshold = 2* and *min.pct = 0.35* for a stringent threshold.

To map the cell clusters to the published oligodendrocyte lineage groups, cell similarity score analysis was conducted using the “TySim” package (version: 0.1.0) (https://github.com/yu-lab-vt/CMC/tree/CMC-TySim).^61^ This analysis provides a quantitative metric of single-cell-to-target-cell-type similarity based on snRNA-seq data and the DEG list of the target cell type (Table S2). It quantifies the degree of similarity between each single cell and the target cell type.

For the gene ontology (GO) term activity analysis and GO term-based clustering analysis, the “GOTermActivity” package (version: 0.3.1) (https://github.com/yu-lab-vt/CMC/tree/CMC-GOTermActivity) was utilized on the current snRNA-seq dataset.^62^ This package converts snRNA-seq data into a matrix of GO term activity scores, representing the activity levels of individual GO terms in each cell. Each activity score is accessed by analyzing the overall expression of gene set associated with the specific GO term, along with the background expression of each gene in each specific cell. This is achieved by jointly modeling the cell-specific and gene-specific factors within the snRNA-seq data.

### Electrophysiology and imaging

To label premyelinating oligodendrocytes, 100 mg/kg of 4-hydroxytamoxifen (4HT, Cayman Chemical, Cat#1730850) was injected to *Enpp6-CreER^T2^; Ai9/+* mice at P14 via intraperitoneal injection. The 4HT injected mice were sacrificed at P16 (2 dpi) and P18 (4 dpi) to examine electrophysiological properties of oligodendrocytes at different differentiation stages. The mice were euthanized with overdoes of isoflurane. The brains were quickly extracted from skull and put in ice chilled artificial cerebrospinal fluid (aCSF, consisting of 125 NaCl, 2.5 KCl, 1.25 NaH_2_PO_4_, 25 Glucose, 25 NaHCO_3_, 1 MgCl_2_, 2 CaCl_2_ in mM, pH 7.4 adjusted with NaOH or HCl). The prefrontal cortex (PFC) of the mouse brain was sectioned with 200 µm thickness in iced aCSF, utilizing the vibratome (Leica, VT1200s). Brain slices containing PFC were recovered in the incubation chamber containing aCSF bubbled with carbogen (95% O_2_ and 5% CO_2_), where they were maintained for 30 min at 34°C and thereafter at room temperature (24°C).

The DsRed positive cells in the slice were visualized using an infrared differential interference contrast microscope (AxoExaminer, Zeiss, Oberkochen Germany) with a 40X water-immersion objective and TRITC filter. Whole-cell patch-clamp recordings were performed in the chamber perfused with aCSF at room temperature (24°C) using an EPC-10 amplifier controlled by PATCHMASTER software (HEKA, Elektronik, Lambrecht/Pfalz, Germany). Voltage clamp recording was performed from DsRed^+^ oligodendrocytes with the pipette filled with solution containing (in mM, pH adjusted to 7.3 with KOH): 130mM K-gluconate, 10mM KCl, 10mM HEPES, 0.2mM EGTA, 4mM MgATP, 0.3mM NaGTP, 5mM Na-phosphocreatine and 0.1 Alexa Fluor™ 488 Hydrazide (Invitrogen, Cat#A10436). Steady-state currents against various voltage steps (−100 mV to −10 mV) were recorded from each cell. The input resistance (R_input_) was calculated by measuring the steady-state current changes from −70 mV to −50 mV of holding membrane potential. After recording, the slices containing the patched cells were fixed with 4% PFA. To examine the colocalization between DsRed and infused intra cellular dye (Alexa fluor 488), the slices were viewed with laser lines at 488 nm and 568 nm using a 40x oil-immersion objective on a confocal laser-scanning microscope (Nikon A1).

### Whisker trimming

For the P5 whisker trimming paradigm, P5 pups were whisker trimmed unilaterally and were repeatedly trimmed daily. Single intraperitoneal 4HT injections were administered at P8, P9, and P10 at a dosage of 100 mg/kg, followed by tissue collections at P14. For the P8 whisker trimming paradigm, P8 pups were administered with 4HT at a dosage of 150 mg/kg while whisker trimming was performed unilaterally. Trimming was repeatedly performed daily up to the age of collection.

